# Clustering-independent estimation of cell abundances in bulk tissues using single-cell RNA-seq data

**DOI:** 10.1101/2023.02.06.527318

**Authors:** Rachael G. Aubin, Javier Montelongo, Robert Hu, Elijah Gunther, Patrick Nicodemus, Pablo G. Camara

## Abstract

Single-cell RNA-sequencing has transformed the study of biological tissues by enabling transcriptomic characterizations of their constituent cell states. Computational methods for gene expression deconvolution use this information to infer the cell composition of related tissues profiled at the bulk level. However, current deconvolution methods are restricted to discrete cell types and have limited power to make inferences about continuous cellular processes like cell differentiation or immune cell activation. We present ConDecon, a clustering-independent method for inferring the likelihood for each cell in a single-cell dataset to be present in a bulk tissue. ConDecon represents an improvement in phenotypic resolution and functionality with respect to regression-based methods. Using ConDecon, we discover the implication of neurodegenerative microglia inflammatory pathways in the mesenchymal transformation of pediatric ependymoma and characterize their spatial trajectories of activation. The generality of this approach enables the deconvolution of other data modalities such as bulk ATAC-seq data.

## Introduction

Biological tissues are complex systems composed of millions of cells interacting to produce biological function. Characterizing the cellular composition and heterogeneity of tissues is thus fundamental to understanding the relation between cellular phenotypes and tissue function, and it has been a major area of investigation for over a century^1-3^. Advances in high-throughput single-cell RNA sequencing (RNA-seq) have revolutionized the study of tissue composition by enabling the transcriptomic characterization of cell types and states without the need for pre-defined markers^4,5^. However, establishing robust associations between tissue cell composition and other data, such as clinical data, requires generating, profiling, and analyzing large cohorts of samples, which is often technically, computationally, and financially prohibitive by single-cell RNA-seq. In addition, the tissue dissociation and cell encapsulation techniques involved in single-cell RNA-seq can lead to the underrepresentation of some cell populations^6^. Since transcriptomic profiling of tissues at the bulk level does not suffer from these limitations, an enticing alternative is to use the bulk-level gene expression profile of each sample to computationally infer the abundance of each cell population in the sample^7,8^. This approach is known as gene expression deconvolution.

Current methods for gene expression deconvolution use matrix regression, such as support vector^9,10^, least-squares^11-15^, elastic net^16^, or least absolute deviation regression^17^, to represent the overall gene expression profile of the tissue as a linear combination of cell type-specific gene expression signatures (Figure 1A). These gene expression signatures are directly provided by the user or built from a reference single-cell RNA-seq dataset by clustering and differential gene expression analysis of the single-cell transcriptomes. These approaches work particularly well when the cell types present in the sample have very distinct gene expression profiles and form discrete clusters in the single-cell gene expression space^18,19^. However, poorly characterized cell states or continuous cellular processes, such as cell differentiation or immune cell activation, cannot be accurately described in terms of discrete cell populations and often involve colinear gene expression signatures. These nuances are usually lost inside broader populations, limiting the resolution to detect small changes in cell state. In addition, by averaging transcriptomic variability within cell clusters, the output of these methods strongly depends on the choice of clustering algorithm and parameters. Consistent with these limitations, two recent studies found that the specificity of the reference gene expression signatures that are used is the greatest determinant of accuracy in current methods for gene expression deconvolution^19,20^. Therefore, there is a need for clustering-independent approaches that can take full advantage of reference single-cell RNA-seq data to infer cell abundances in bulk tissue samples with high phenotypic resolution.

**Figure 1.**
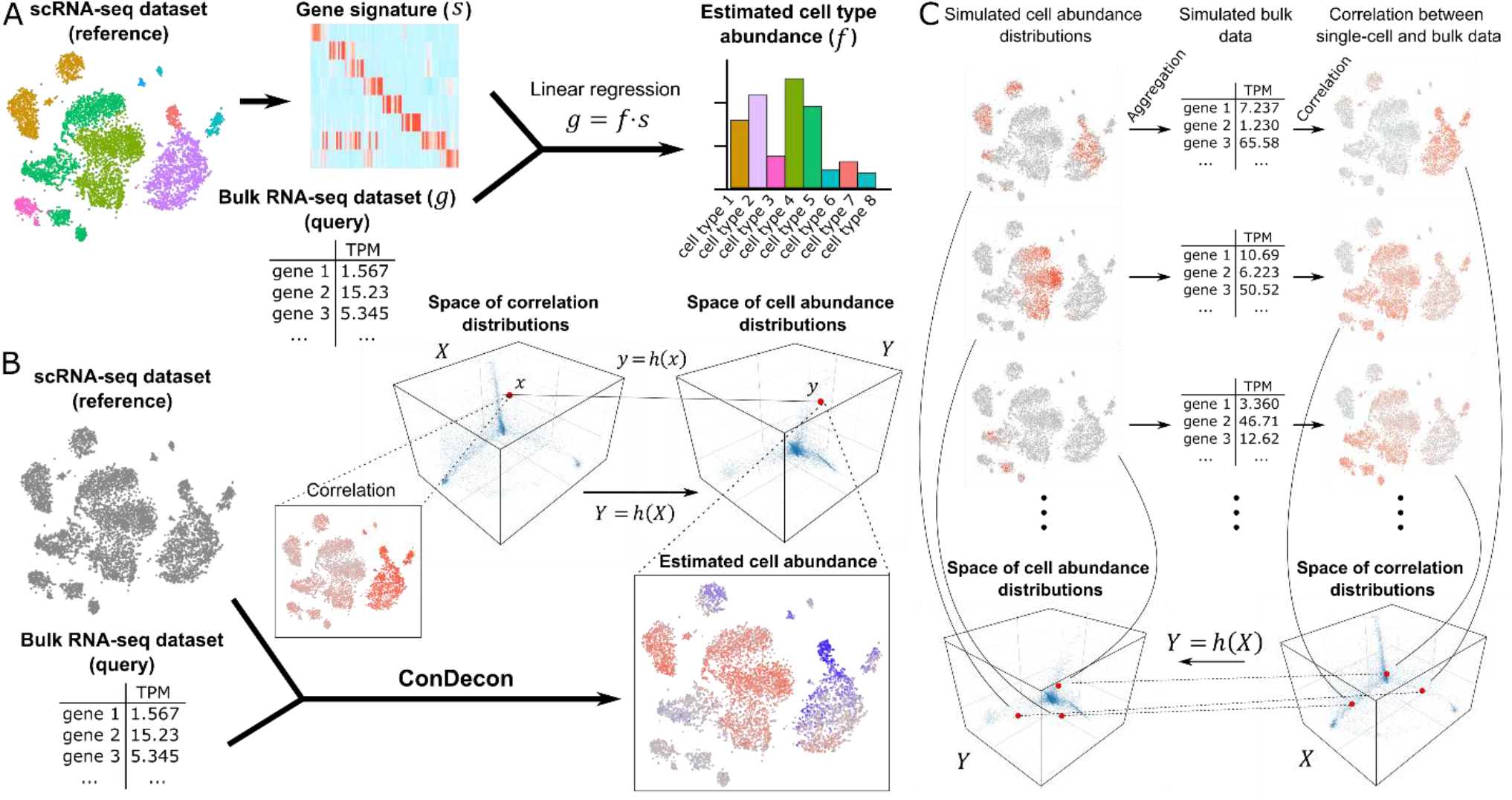
A clustering-independent approach for cell abundance inference from gene expression data of bulk tissues. **(A)** Conventional methods for gene expression deconvolution of bulk tissues cluster a reference single-cell RNA-seq dataset into discrete cell populations and perform differential gene expression to build a gene expression signature matrix for the discrete cell populations. The problem of estimating discrete cell type abundances is then formulated as a linear regression problem. **(B)** The approach of ConDecon to gene expression deconvolution differs substantially from that of conventional methods. It takes as input a bulk RNA-seq query dataset and a reference single-cell RNA-seq dataset. It then computes the rank correlation between the gene expression profiles of the bulk RNA-seq dataset and each cell in the single-cell dataset using the most variable genes. The resulting correlations are represented by a point in the space of possible correlation distributions with support on the single-cell RNA-seq latent space. ConDecon then maps that point into a point in the space of possible cell abundance distributions with support on the single-cell RNA-seq latent space. **(C)** The model of ConDecon is trained by simulating multiple cell abundance distributions with support on the single-cell RNA-seq latent space by means of a Gaussian mixture model. For each simulated distribution, a synthetic bulk RNA-seq dataset is constructed by aggregating the gene counts, and the rank correlation between the gene expression profiles of the synthetic bulk RNA-seq dataset and each cell in the single-cell dataset is computed using the most variable genes. The paired cell abundance and correlation distributions are then used to learn the function *h* that maps the spaces of possible correlation distributions and cell abundance distributions with support on the single-cell RNA-seq latent space. See also Figure S1.

Here we present a deconvolution method, named ConDecon, for inferring cell abundances from gene expression data of bulk tissues without relying on cluster labels or cell-type specific gene expression signatures at any step. The aim of ConDecon is to infer a probability distribution across a reference single-cell RNA-seq dataset that represents the likelihood for each cell in the reference dataset to be present in the query bulk tissue. Through multiple analyses of simulated and real data from well-characterized systems with known ground truth, we demonstrate that ConDecon can be used to accurately estimate cell abundances in bulk tissues composed of discrete cell types and continuous cellular processes, where the application of current deconvolution methods is limited. The estimates that result from aggregating ConDecon’s cell probabilities across cells of the same type are highly concordant with flow cytometry measurements, mirroring state-of-the-art clustering-based deconvolution methods.

We demonstrate the utility of ConDecon to uncover new biological insights by applying it to single-cell, bulk, and spot-based spatial transcriptomic data of pediatric ependymal tumors^21,22^. Through these analyses, we discover consistent changes in the expression program of tumor-infiltrating microglia associated with the mesenchymal transformation of tumor stem cells. By mapping the continuous differentiation trajectories of microglia and tumor cells from the reference single-cell RNA-seq dataset onto the spatial transcriptomic data, we identify distinct spatial patterns of microglia activation around mesenchymal tumor regions. We find that microglia in these areas develop a phenotype akin to that in Parkinson’s and Alzheimer’s disease lesions, marked by increased expression of *GPNMB*^23-25^. In addition, we highlight the broad applicability of the approach implemented in ConDecon to other omics data modalities, including the estimation of cell abundances in bulk ATAC-seq data using reference single-cell ATAC-seq data.

Altogether, these applications demonstrate that ConDecon enables previously elusive analyses of dynamic cellular processes in bulk tissues and represents an increase in functionality and phenotypic resolution with respect to current methods for gene expression deconvolution.

### Design

To overcome the inherent limitations of cell-type specific gene expression signatures in the deconvolution of gene expression data from bulk tissues, we developed ConDecon, a clustering-independent method for inferring changes in cell abundance based on reference single-cell RNA-seq data provided by the user. ConDecon uses the gene expression count matrix and latent space of the reference single-cell RNA-seq dataset to estimate the likelihood of each cell in the dataset to be present in the query bulk tissue sample (Figure 1B). For that purpose, it assumes that the similarity between the gene expression profile of cells in the single-cell dataset and that of the bulk tissue sample, as measured by their rank correlation, can be used as a proxy for inferring this likelihood function (Figure 1B, Methods). The use of correlations between gene ranks instead of gene expression values is motivated by their greater stability against the technical differences between single-cell, single-nucleus, and bulk gene expression measurements (Figure S1A). A similar strategy has been recently proposed for identifying cell populations from single-cell data associated with a given phenotype^26^. The goal of ConDecon is to learn a map *h*(*X*): *X* → *Y* between the space *X* of possible rank correlation distributions and the space *Y* of possible probability distributions on the single-cell gene expression latent space (Box 1, Methods). To that end, it introduces coordinates in *X* and *Y* by expanding the distributions in a basis, such as principal components or diffusion maps, and represents *h*(*X*) as a polynomial function on the coordinates.

To learn *h*(*X*), ConDecon simulates bulk transcriptomic data by aggregating the gene expression profiles of cells sampled from the single-cell reference dataset according to a randomly generated mixture of Gaussian distributions with a variable number of components (Figure 1C). The use of smoothly varying probability distributions to train ConDecon contributes to the regularization of the output (Methods). Each simulated probability distribution represents a point *y* ∈ *Y*. The rank correlation coefficients between the gene expression profiles of the simulated bulk dataset and each of the cells in the single-cell dataset then provide a point *x* ∈ *X* such that *y = h*(*x*) (Figure 1C). By using this procedure to simulate many bulk datasets, it is possible to fit the model for *h*(*X*). With the fitted model, ConDecon can then infer the distribution of cell abundances for any query bulk sample of the same tissue type as the single-cell reference dataset. This clustering-independent approach is therefore conceptually different from the regression-based approach used by current methods for gene expression deconvolution and takes full advantage of all the variability contained in the reference single-cell dataset.

We have implemented ConDecon as an open-source package in R. The Detailed Protocol provided as a Supplemental Item to this article demonstrates step-by-step the application of ConDecon to different data modalities.

## Results

### Estimation of cell abundances in simulated RNA-seq data of discrete cell types and continuous cellular processes

To demonstrate the ability of ConDecon to infer changes in cell state, we simulated single-cell RNA-seq data from a broad range of configurations using the algorithm Splatter^27^. Splatter uses a gamma-Poisson model to simulate gene-by-cell RNA count matrices of complex tissues. From each simulated single-cell dataset, we derived several query bulk RNA-seq datasets by non-uniformly sampling cells from the single-cell dataset and aggregating their gene expression profile. We then used ConDecon to estimate cell abundances in each simulated bulk dataset and compared them to the ground-truth abundances.

We first tested ConDecon in simple scenarios where the bulk tissue consists of discrete cell populations. In these simulations, each cell population was taken to be homogeneous up to some random variability. We generated 27 single-cell RNA-seq datasets consisting of 3, 6, or 9 cell types and varying degrees of differential gene expression. From each dataset, we derived 25 bulk RNA-seq datasets where we varied the sampling probability of each cell type to simulate different cell type abundances. To quantify the concordance between the estimated and simulated cell abundances, we aggregated the estimates of ConDecon across cells of the same type. The estimated and simulated abundances for each cell type were strongly correlated in most cases (Figures 2A, 2B, and S2A, average Pearson’s *r =* 0.75, average *p*-value *=* 0.06). This correlation was higher for samples with fewer cell types or higher differential gene expression (Figure 2B). Thus, ConDecon can be efficiently used to deconvolve gene expression in bulk tissues consisting of discrete cell populations, where standard methods for gene expression deconvolution can also be applied.

**Figure 2.**
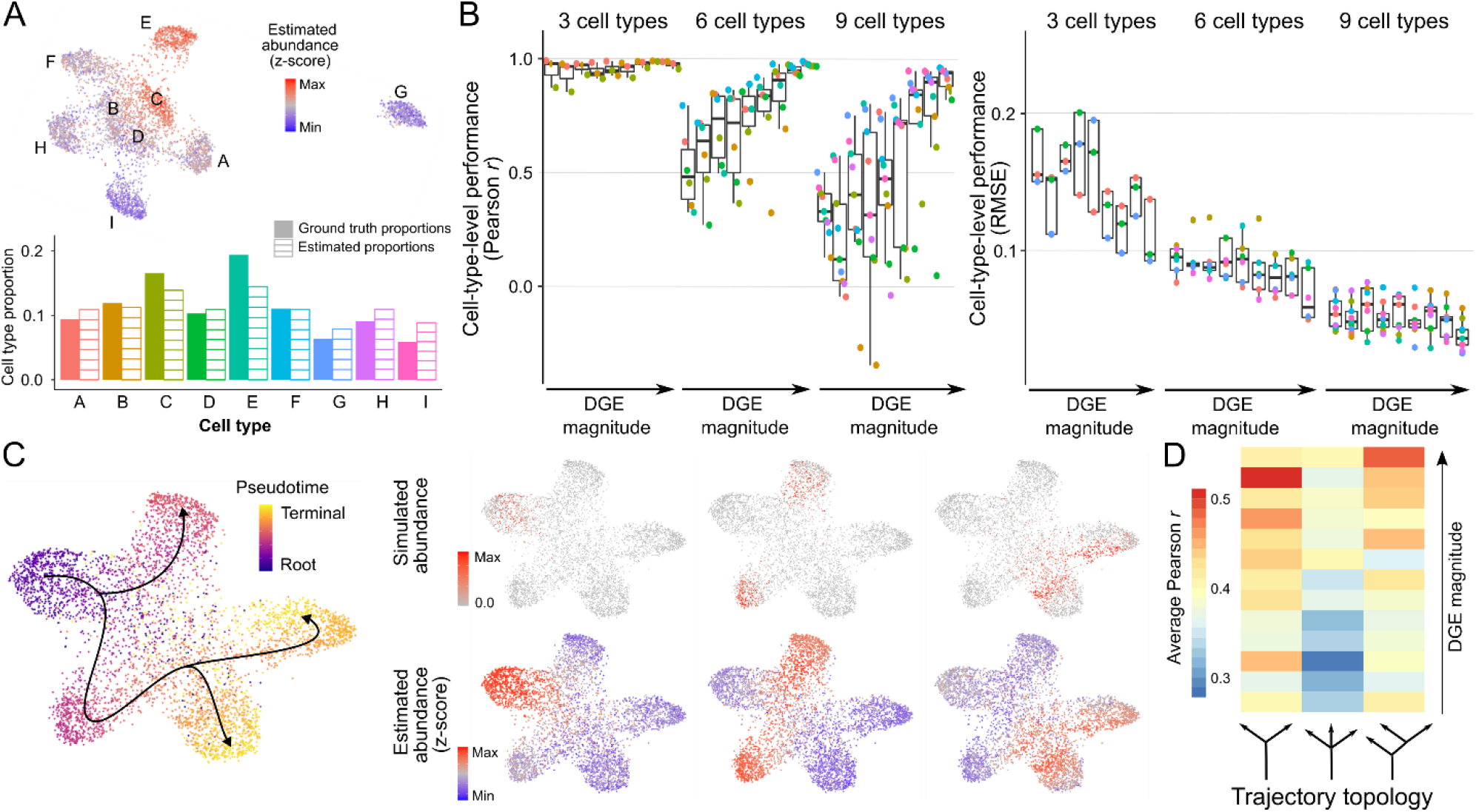
Estimation of cell abundances from simulated gene expression data of discrete cell populations and continuous cellular processes. **(A)** Estimation of cell population abundances in a simulated bulk RNA-seq dataset with 9 discrete cell populations (named A to I). Single-cell RNA-seq data was simulated using Splatter and the gene expression profiles of individual cells sampled with different probability from each cell population were pooled to construct a synthetic bulk RNA-seq dataset. The UMAP representation of the single-cell RNA-seq data is colored by ConDecon’s estimated cell abundances (top). The aggregated cell abundance estimates across each cell population recapitulate the simulated cell population abundances (bottom). **(B)** Pearson correlation coefficient and root mean square error (RMSE) between the simulated and estimated abundances for each cell population in 675 simulated bulk RNA-seq datasets with 3, 6, or 9 discrete cell populations and varying degree of differential gene expression (DGE). Average Pearson’s correlation coefficient *r =* 0.75, average *p*-value *=* 0.06. **(C)** Cell abundance estimation in 3 out of 25 simulated bulk RNA-seq datasets of a continuous cell differentiation process with 1 precursor and 3 terminally differentiated cell states. The UMAP representation of the simulated single-cell RNA-seq data is colored by the simulated pseudotime (left). The cell abundances estimated with ConDecon in each of the simulated bulk RNA-seq datasets (right, bottom) recapitulate the simulated abundances that were used to construct the bulk data (right, top). Cell abundances have been standardized across samples. **(D)** Average Pearson correlation coefficient between the simulated and estimated cell abundances in 975 simulated bulk RNA-seq datasets with 3 topologies for the cell differentiation trajectories and varying degree of differential gene expression (DGE). Average Pearson’s correlation coefficient *r =* 0.40, average *p*-value *=* 2 × 10^−9^. See also Figure S2.

We next simulated single-cell RNA-seq data from continuous cellular processes, such as cell differentiation, for which the application of conventional gene expression deconvolution methods is contrived. We generated 39 single-cell RNA-seq datasets consisting of cell differentiation trajectories with three different topologies and varying differential gene expression levels. Each trajectory consisted of a precursor and two or three terminally differentiated cell states (Figure 2B). For each simulated single-cell RNA-seq dataset, we generated 25 bulk datasets by sampling cells based on a random Gaussian kernel on pseudotime to simulate the asynchrony of the cell differentiation process (Figure 2C). The single-cell abundance estimates of ConDecon were again strongly correlated with the ground-truth abundances (Figures 2C, 2D, and S2D, average Pearson’s *r =* 0.40, average *p*-value *=* 2 × 10^−9^) and their accuracy improved with the amount of differential gene expression (Figure 2D). Since estimating individual cell abundances is a more challenging problem than estimating cell population abundances, a lower correlation coefficient was observed in this case.

To evaluate the stability of these results, we repeated these analyses using different choices for the parameters of ConDecon, including the dimensionality of the spaces *X* and *Y* and the number of variable genes used in computing the rank correlation. This analysis showed that the estimates of ConDecon are stable against different choices for its parameters, only observing a substantial decrease in the accuracy of the estimates for small values of the parameters (less than 500 genes and 5 dimensions) (Figures S2B, S2C, and S2E).

Taken together, the application of ConDecon to simulated data demonstrates the validity of its clustering-independent approach to deconvolve gene expression data from complex tissues consisting of both discrete cell populations and continuous cellular processes.

### Stability of cell abundance estimates in real RNA-seq data

We next evaluated the stability of the cell abundance estimates produced by ConDecon using real data. For that purpose, we considered two published bulk RNA-seq datasets consisting of 8 bone marrow^28^ and 12 peripheral blood mononuclear cell (PBMC)^10^ samples for which paired fluorescence-activated cell sorting (FACS) data were available. We applied ConDecon to the two datasets using a broad range of parameters (Methods). For each run, we aggregated the resulting single-cell probabilities across each cell type to infer cell type abundances, which we then compared to the FACS data using Pearson’s correlation and root mean square error (RMSE) across samples (cell-type-level performance) or cell types (sample-level performance). Consistent with the results of our simulations, we found that a minimum of approximately 500 variables genes and 5 dimensions were needed for the cell type abundance estimates to be in good agreement with the FACS data (Figures S3A and S3B). We also tested whether there was any advantage in modeling *h*(*x*) as a quadratic polynomial on the coordinates instead of as a linear function. Since the number of coefficients to be fitted in a quadratic polynomial is substantially larger ((*D*^3^ + 3*D*^2^)/2 instead of *D*^2^, with *D* the number of dimensions of the latent space), we increased the size of the training dataset by one order of magnitude to ensure an adequate fit. In this analysis, we did not observe a substantial improvement in the accuracy of the results by using a quadratic polynomial (Figures S3A and S3B). This is consistent with the approximate linearity of *h*(*x*) observed in our simulations (Box 1 and Figures S1B and S1C). Thus, due to the added computational cost of generating larger training datasets, we decided to model *h*(*x*) as a linear function in subsequent analyses.

A potential limitation of using gene rank correlations as the basis for gene expression deconvolution is that the inferred cell abundances are not unique (Box 1, Methods). However, this concern can be safely disregarded in standard single-cell datasets since the uncertainty in the estimates for datasets consisting of hundreds of variable genes is expected to be smaller than other sources of variability. To verify this, we repeated the analyses of the bone marrow and PBMC datasets using 20 different random initializations of ConDecon. As expected, the variability of the inferred abundances across runs was significantly smaller than the variability across samples for most cell types (Figure S3C). This demonstrates the effectiveness of ConDecon in inferring cell abundance differences between samples, even in challenging cases like these, where the bulk samples were derived from healthy donors and the observed variability corresponds to small, naturally occurring interindividual variation.

We finally evaluated the characteristics of the reference single-cell RNA-seq data that are needed to obtain accurate cell abundance estimates. For that purpose, we randomly down-sampled the bone marrow and PBMC single-cell data to 25% and 10% of the cells (corresponding to approximately 2,000 and 800 cells, respectively). We found that the cell-type-level and sample-level performance of ConDecon decreased when 10% of the cells were included in the reference single-cell data (Figures S3A and S3B). Moreover, in the case of PBMC data, the cell-type performance was also reduced when considering 25% of the cells (Figure S3A). Based on these observations, we suggest using ConDecon with single-cell datasets consisting of at least 5,000 cells in total and 100 cells per cell type/state.

To assess the impact of large differences in cell type proportions between the reference single-cell and query datasets, we sampled cells from the original single-cell PBMC and bone marrow datasets to construct new single-cell datasets where all cell types were equally represented. To avoid sampling with repetition, the size of these datasets was limited to 700 and 1,630 cells, respectively. In addition, we generated single-cell datasets of the same size and cell type proportions as the original single-cell dataset and the FACS data. Through these analyses, we found that the cell-type-level performance of ConDecon increased when all cell types were equally represented in the reference dataset (Figures S3A and S3B). However, the shift in cell type abundances in the reference data negatively impacted the sample-level performance.

Hence, while ConDecon’s inferences of cell type abundance variation across samples are robust against large differences between the reference and query datasets, achieving accurate estimates of the relative abundances within each sample requires single-cell RNA-seq datasets from the same tissue type as the bulk data.

We also evaluated the effect of large mismatches in the cell types present in the reference and query datasets. We considered bone marrow bulk RNA-seq data and bone marrow and kidney single-cell RNA-seq data from the Tabula Muris Senis^29^. The single-cell datasets contained multiple cell types exclusive to the kidney (e.g., proximal tubule cells) or the bone marrow (e.g., myeloid progenitors) (Figure S3D). We combined the two single-cell datasets into a single reference to deconvolve the bone marrow bulk RNA-seq data. Consistent with our previous observations, the inferences of ConDecon were affected by the large mismatch between the reference and query datasets, and 24% of the probability mass was assigned to kidney-specific cell populations on average (Figure S3D). In this regard, we noted that the distance between the query data point and the nearest training data points in the space of probability distributions *Y*, normalized by the average distance between training data points, is a good indicator of the quality of the inferences made by ConDecon, with smaller distances corresponding to more accurate inferences (Figure S3D).

Altogether, these analyses indicate that while ConDecon is particularly suited for inferring subtle variations in cell abundances across samples and is robust against a broad range of parameters, inferring accurate relative abundances within each sample necessitates reference datasets from the same tissue type as the bulk samples. However, we do not expect this to be a major limitation, given the current broad availability of single-cell data from most tissues^30,31^.

### The accuracy of ConDecon’s estimates of discrete cell type abundances mirrors that of state-of-the-art clustering-based methods

We used a published benchmarking pipeline^32^ to systematically compare the cell type abundance estimates of ConDecon with those produced by 17 other methods for gene expression deconvolution. The pipeline uses single-cell RNA-seq data to simulate bulk RNA-seq datasets containing mixtures of discrete cell types. It then evaluates the accuracy and stability of the estimates produced by each algorithm when none or one cell type in the query sample is absent in the reference data^32^. For these comparisons, we used five single-cell RNA-seq datasets of PBMCs^10^, pancreas^11,33^, bone marrow^28^, and kidney^34^. Each algorithm was evaluated based on Pearson’s and Lin’s concordance correlation coefficients, as well as the root mean squared error (RMSE), between the estimated and simulated cell abundances across samples and cell types^32^. In agreement with previous comparative studies of gene expression deconvolution^19,20,32^, no single method performed best across all datasets and metrics.

Nonetheless, ConDecon ranked among the best-performing methods according to several metrics (Figures 3A and S4). Its cell type abundance estimates had an average combined Pearson’s correlation of 0.91 with the ground truth, surpassing 16 of the 17 other methods according to this metric. While its performance based on Lin’s concordance correlation was lower, it remained a strong performer by this metric as well, with an average combined Lin’s correlation only 7% lower than the top-performing method (Figures 3A and S4).

**Figure 3.**
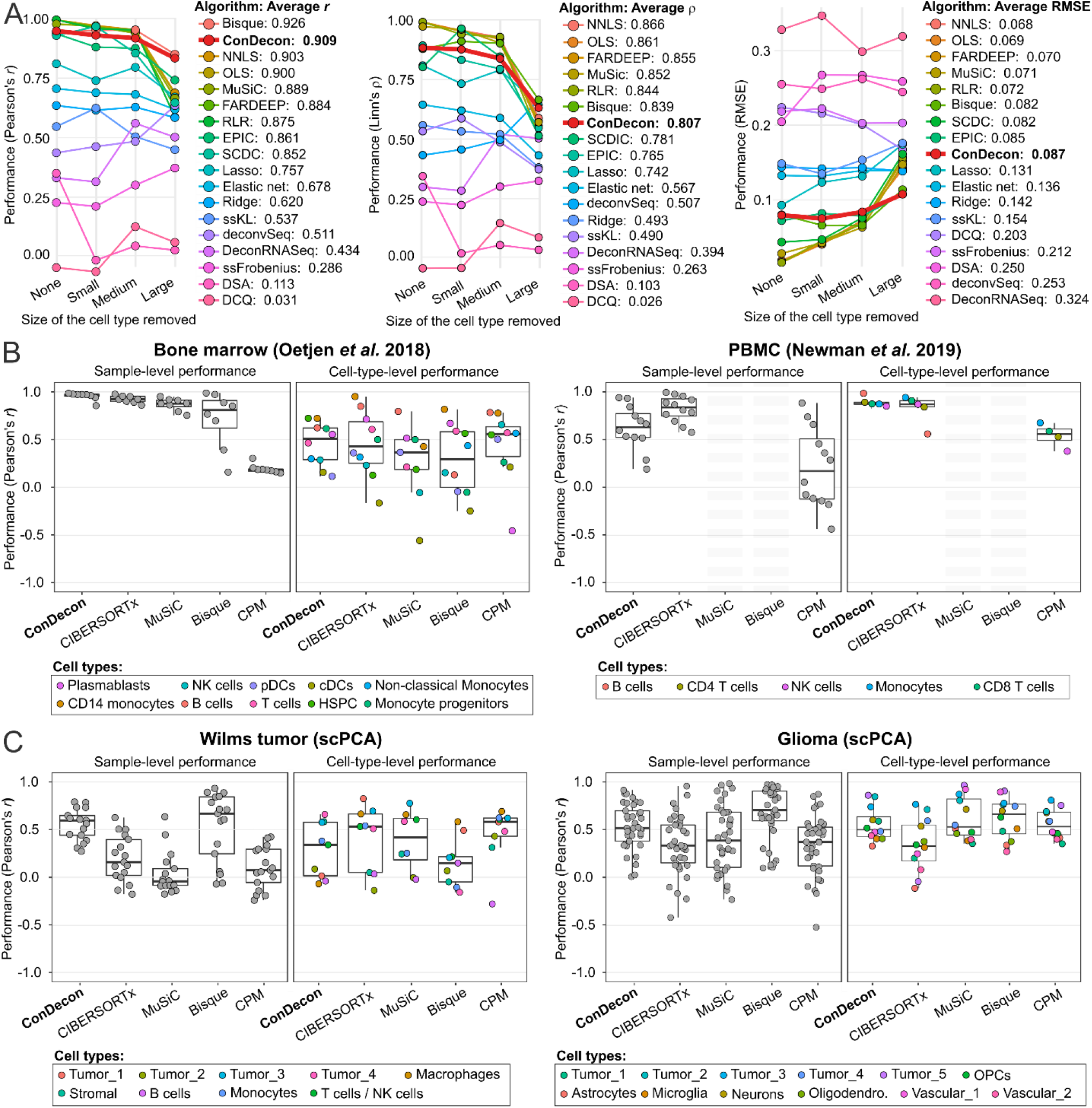
Benchmarking the cell type abundance estimates of ConDecon in comparison to current gene expression deconvolution methods. **(A)** The cell abundance estimates of ConDecon were aggregated into cell type abundance estimates and compared with those of 17 other deconvolution methods across 5 datasets using the benchmarking pipeline of Avila-Cobos *et al*.^32^. For each algorithm, the Pearson’s correlation coefficient (left), the Lin’s concordance correlation coefficient (center), and the root mean squared error (RMSE) (right) of the estimates, combined across samples and cell types, is shown for cases where there is none, one small, one medium, or one large cell population missing in the reference single-cell data. **(B, C)** Comparison between cell type abundance estimates derived from FACS and single-nucleus RNA-seq data and those from ConDecon and 4 other deconvolution methods specifically devised to use single-cell RNA-seq data as reference data. Two bulk RNA-seq datasets consisting of 8 bone marrow^28^ (B, left) and 12 PBMC^10^ (B, right) samples, for which paired FACS data are available, as well as two bulk RNA-seq datasets consisting of 17 Wilms tumor (C, right) and 37 pediatric glioma (C, left) samples, for which paired single-nuclei RNA-seq data are available, were considered for this evaluation. The sample-level and cell-type-level Pearson’s correlation coefficient are shown for each algorithm in each dataset. We were unable to apply MuSiC and Bisque to the PBMC dataset since these methods require that the reference single-cell RNA-seq data consists of at least 2 biological replicates. See also Figures S4 and S5.

As expected, the accuracy of the estimates produced by all algorithms decreased as the size of the missing population in the reference data increased. However, ConDecon and Bisque^35^ were the most robust against missing large cell populations (Figures 3A and S4). In terms of specific tissues, the estimates of ConDecon were particularly accurate in the kidney and bone marrow datasets, where it outperformed the other 17 methods according to two out of three metrics (Figure S4). In contrast, its performance in the two pancreas datasets was moderate.

A caveat of the benchmarking pipeline is that the bulk RNA-seq data are simulated from single-cell RNA-seq data and may lack some of the technical features present in actual bulk RNA-seq datasets. To address this limitation, we considered the two aforementioned PBMC^10^ and bone marrow^28^ bulk RNA-seq datasets with paired FACS data from the stability analysis, as well as two bulk RNA-seq datasets from the single-cell pediatric cancer atlas (scPCA)^36^, consisting of 17 Wilms tumor and 37 pediatric glioma samples for which paired single-nucleus RNA-seq (snRNA-seq) data were available. We compared the estimates of ConDecon in the four bulk RNA-seq datasets with those of CIBERSORTx^10^, MuSiC^12^, Bisque^35^, and CPM^37^, as well as with the FACS or snRNA-seq cell type abundances. Like ConDecon, these gene expression deconvolution methods have been specifically devised to use single-cell RNA-seq data as a reference. Specifically, CIBERSORTx accounts for platform-specific variation when comparing single-cell and bulk gene expression levels, while MuSiC and Bisque leverage multi-subject single-cell expression data to improve the accuracy of the estimates. These three algorithms seek to infer the abundance of each cell type in the query sample, whereas CPM aims to reconstruct the continuous spectrum of cell states within a single query cell type. For that purpose, CPM partitions the gene expression space of the cell type into smaller discrete domains and uses a bootstrapped support vector regression approach to infer the abundance of each domain^37^.

Consistent with our results from the benchmarking pipeline, no single method outperformed the others across all datasets and metrics (Figures 3B, 3C, and S5). However, when it came to estimating relative variations in cell abundances across cell types (sample-level performance), ConDecon consistently exhibited the highest or second-highest performance in each of the four datasets according to all metrics. Specifically, ConDecon and CIBERSORTx provided the best sample-level performance in the two datasets with FACS data, while ConDecon and Bisque provided the best sample-level performance in the two solid tumor datasets. We were unable to apply MuSiC and Bisque to the PBMC dataset, as these methods require the reference single-cell RNA-seq data to have at least two replicates.

Regarding the estimation of relative abundances across samples (cell-type-level performance), all methods exhibited relatively poor performance in the four datasets, with Lin’s concordance correlation usually falling below 0.5 (Figures 3B, S3C, and S5). Nonetheless, the cell-type-level performance of ConDecon remained the best or second-best according to all metrics in each of the two datasets with FACS data, while its cell-type-level RMSE was the smallest in three of the four datasets. In contrast, the cell-type-level Lin’s and Pearson’s correlation coefficients of ConDecon in the two solid tumor datasets were lower compared to those of other methods.

Altogether, these results demonstrate that aggregating the single-cell abundance estimates of ConDecon into discrete cell type abundances yields estimates that exhibit comparable accuracy to those produced by state-of-the-art deconvolution methods. In particular, the clustering-independent approach implemented in ConDecon does not disadvantage the estimation of discrete cell type abundances compared to current clustering-based approaches, while it enables a higher phenotypic resolution in the study of continuous cellular processes, as we show below.

### Inference of continuous changes in B-cell maturation with ConDecon

Having tested ConDecon with tissues that consist of cell types with very distinct gene expression profiles, we next used it to study changes in single-cell abundance associated with continuous cellular processes. We considered single-cell and bulk RNA-seq data of bone marrow from mice aged between 1 and 27 months^29^ and used these data to study changes in cell abundance associated with development and aging. We used the well-characterized changes in B-cell abundance that occur during postnatal development^38^ as a test system. Using an integrated representation of these single-cell data with no age labels as a reference, ConDecon was able to recapitulate from the bulk data the continuous transition from an abundance of pro B-cells in young mice (≤ 3 months) to an abundance of naïve mature B-cells in fully developed mice (Figures 4 and S6A, Pearson’s correlation between the age of mice and the average inferred pseudotime of the B-cells in each mouse *r* = 0.77, *p*-value = 2 × 10^−11^), in agreement with previous results based on FACS data^38^. In addition, ConDecon predicted changes associated with aging in other bone marrow cell populations, such as an increase of megakaryocyte-erythroid progenitors and NK cells with age^39,40^ (Spearman’s *ρ* = 0.88 and 0.81, *p*-value = 0.002 and 0.008, respectively). Compared to the alternative approach of sub-clustering the continuous B-cell trajectory into discrete cell subpopulations and using conventional deconvolution methods to infer the abundance of each subpopulation, ConDecon showed a higher power to identify changes in cell abundance (Figure 4E), possibly because conventional methods do not account for intra-cluster variability in cell state abundance. For instance, although the overall immature B-cell population was not enriched in 1-month-old mice, a subset of these cells with a gene expression profile close to that of precursor B-cells was already present at this age (Figure 4B). Thus, ConDecon’s inferences are not restricted to discrete cell populations and can be used to infer changes in cell abundance along continuous cellular trajectories with high resolution.

**Figure 4.**
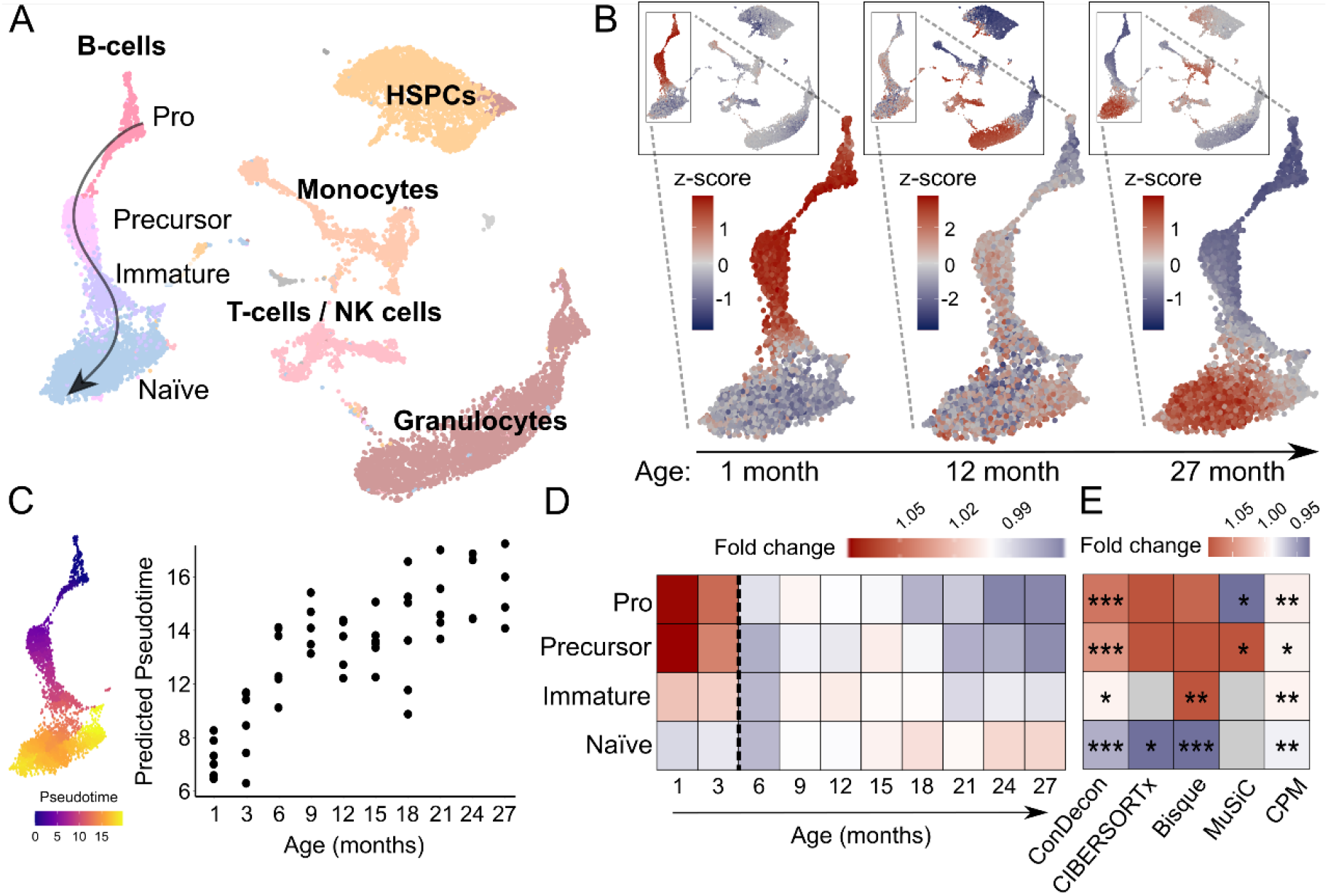
Identification of age-associated changes in B-cell maturation using bulk bone marrow tissues. **(A)** UMAP representation of the single-cell RNA-seq data of 14,107 cells from the bone marrow of 13 mice with ages between 1 and 27 months profiled by the Tabula Muris Consortium^29^. The representation is labelled by the major annotated cell populations. The developmental lineage of B-cells has been subclustered into Pro B-cells, precursor B-cells, immature B-cells, and mature naïve B-cells. HSPCs: hematopoietic stem and progenitor cells. Single-cell abundances inferred by ConDecon for three bone marrow samples from 1-, 12-, and 27-months old mice profiled with bulk RNA-seq. The youngest mouse has a high abundance of Pro and precursor B-cells, whereas the oldest mouse has a high abundance of mature B-cells. Cell abundances have been standardized across mice to visualize variability. Average pseudotime inferred by ConDecon for the B cells in each bulk sample as a function of the mice age, for bone marrow samples of 53 mice profiled with bulk RNA-seq. As expected, the inferred average pseudotime increases with the age of mice. For reference, the UMAP representation of the B-cell lineage colored by the pseudotime is also shown. Pearson’s correlation coefficient *r* = 0.77, *p*-value = 2 × 10^−11^. **(D)** Fold change in the median aggregated cell abundances inferred by ConDecon for each B-cell subpopulation as a function of the age of mice. **(E)** Fold change in the median aggregated cell abundances between young (≤ 3 months) and adult (> 3 months) mice according to the estimates of ConDecon and four other algorithms specifically devised to use single-cell data as reference. Clustering-based methods (CIBERSORTx, Bisque, and MuSiC) had limited power to capture differences between young and adult mice, whereas CPM inferred very small changes in abundance. (2-sided Wilcoxon rank sum test; *: *p*-value ≤ 0.05, **: *p*-value ≤ 0.01, ***: *p*-value ≤ 0.001). See also Figure S6A.

We also compared ConDecon to MeDuSA^41^, a recently developed method for deconvolving abundances from cell differentiation trajectories. While MeDuSA is limited to inferring cell abundances from one-dimensional trajectories pre-specified by the user and is based on partitioning the trajectories into discrete clusters, in cases like the analysis of B-cell maturation presented here, it can provide similar functionalities to those of ConDecon. To perform the comparison, we repeated the analysis of B-cell maturation using MeDuSA, where the cell type labels and B-cell trajectory pseudotime were provided as input in this case. However, although the results of MeDuSA were consistent with those of ConDecon, the inferred pseudotime of the B-cells in each mouse showed a substantially lower correlation with the age of mice (Figure S6B, Pearson’s r = 0.35, p-value = 0.01).

### Clustering-independent estimation of cell abundances using other omics data modalities

The general approach of ConDecon for estimating cell abundances can be applied to other omics data modalities such as spatial transcriptomics or chromatin accessibility data. To evaluate the utility of using ConDecon to deconvolve spot-based spatial transcriptomic data using single-cell RNA-seq data as reference, we considered published Stereo-seq data of 10 zebrafish embryos profiled 3.3 hours post-fertilization (hpf)^42^. At this stage of development, the embryo consists of ∼4,000 blastomere cells arranged in >11 tiers with varying levels of cell differentiation. We used ConDecon to infer the distribution of cell abundances across each tissue section, where each pixel was treated as a bulk sample. Since pixel size in the processed Stereo-seq data is approximately 10 μm^42^, each pixel is expected to receive contributions from 1 to 3 cells. As a reference dataset, we considered single-cell RNA-seq data of 3.3 hpf embryos from the same study and used diffusion pseudotime^43^ to parameterize the differentiation of blastomere cells in these data (Figure 5A). We then used the cell probabilities inferred by ConDecon for each pixel to deconvolve pseudotime and derive trajectories of cell differentiation in the spatial data (Figures 5B, 5C, and S6C). The resulting trajectories recapitulated the known spatial patterns of cell differentiation in the blastodisc, where the differentiation sequence progresses from marginal blastomere cells into deep and superficial blastomere cells^44^ (Figure 5D). Compared to most of the current methods for deconvolving spot-based transcriptomic data^45-49^, the clustering-independent approach of ConDecon can be used to deconvolve continuous features like cell differentiation pseudotime and study the relation between cell differentiation and tissue architecture.

**Figure 5.**
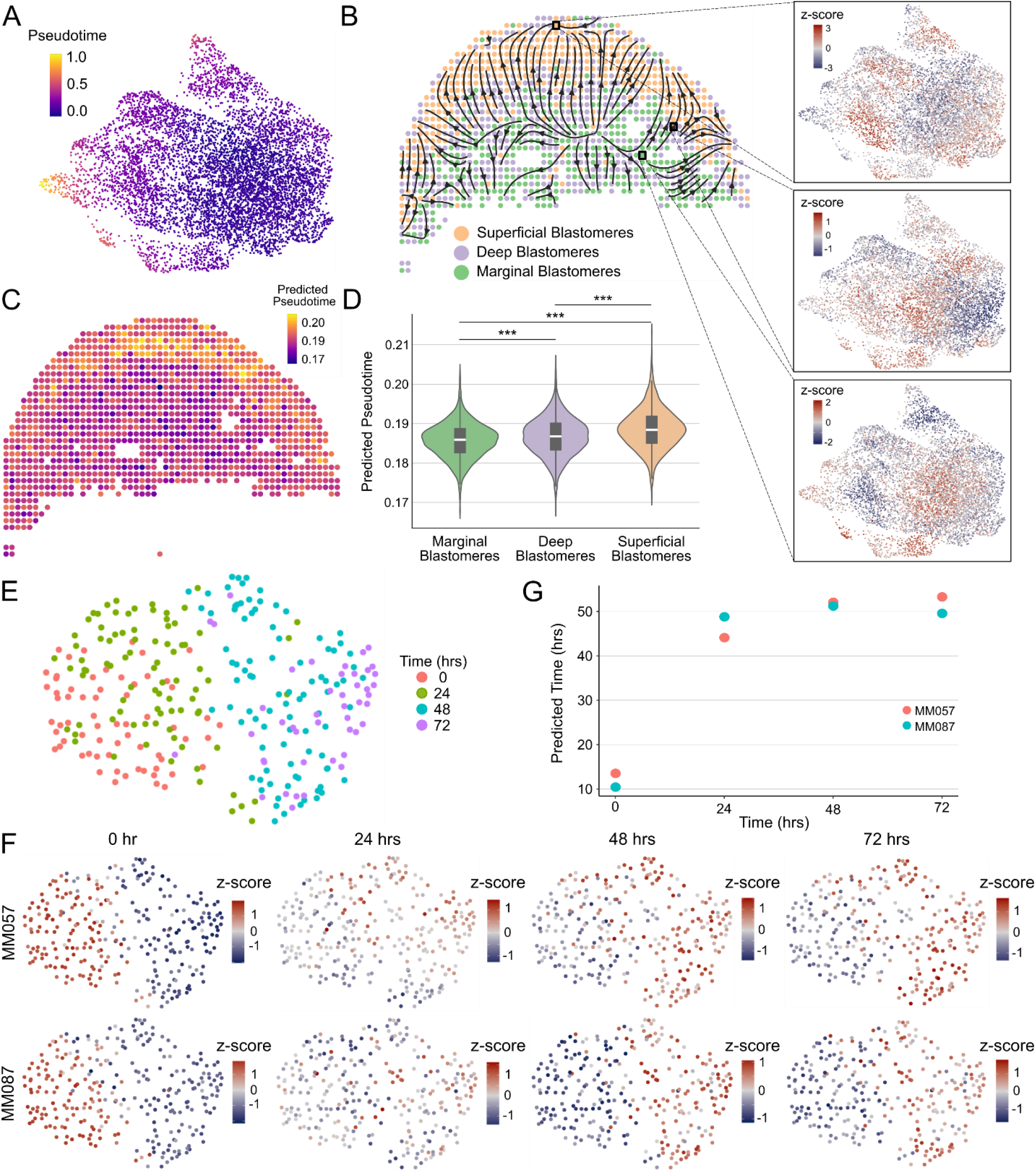
Deconvolution of spatial transcriptomics and ATAC-seq data using ConDecon. **(A)** UMAP representation of the single-cell RNA-seq data of 7,424 blastomere cells from 3.3 hpf zebrafish embryos^42^. The representation is colored by pseudotime associated with the maturation of blastomere cells. **(B)** Spatial representation of a 3.3 hpf zebrafish embryo tissue section profiled with Stereo-seq^42^. Each pixel is labeled according to its majority abundance of marginal, deep, or superficial blastomeres. The spatial cell differentiation trajectories inferred with ConDecon are overlaid on the representation. Differentiation proceeds from marginal blastomeres to deep and superficial blastomeres. The UMAP representation of the reference single-cell dataset colored by the cell abundances estimated with ConDecon is also shown for 3 representative Stereo-seq pixels. Cell abundances have been standardized across samples to highlight variability. **(C)** The same tissue section as in (A) is colored by the average pseudotime of the cells in each pixel estimated with ConDecon. **(D)** Violin plot showing the distribution of estimated pseudotimes for pixels classified as marginal, deep, or superficial across 10 tissue sections profiled with Stereo-seq. Boxes represent the median and interquartile range. Wilcoxon rank-sum test. ***: p-value < 10^−15^. **(E)** UMAP representation of the single-cell ATAC-seq data of a patient-derived melanoma cell line (MM087) profiled 0, 24, 48, and 72 hours after knocking out SOX10^50^. **(F)** The same representation as in (E) is colored by the estimated single-cell abundances for 8 samples from 2 melanoma cell lines (MM057 and MM087) profiled with bulk ATAC-seq 0, 24, 48, and 72 hours after knocking out SOX10. Cell abundances have been standardized across samples to highlight variability. **(G)** Average sampling time estimated with ConDecon for the cells in each of the 8 bulk ATAC-seq samples as a function of the actual sampling time. As expected, the inferred average sampling time for the cells increases with the actual sampling time of the bulk sample. Pearson’s correlation coefficient *r* = 0.83, *p*-value = 0.01. See also Figure S6.

We next evaluated the ability of ConDecon to estimate cell abundances from bulk ATAC-seq data using single-cell ATAC-seq data as a reference. We considered published bulk and single-cell ATAC-seq data of two short-term cultures derived from melanoma patient biopsies^50^. In these cultures, the transcription factor SOX10 was knocked down by siRNA, and cells were sampled at 0, 24, 48, and 72 hours after SOX10 knockdown. To assess the performance of ConDecon in deconvolving bulk ATAC-seq data, we combined the single-cell ATAC-seq data from different time points into a single reference single-cell dataset and compared the sampling time of the cells inferred by ConDecon for each of 8 bulk samples with the actual sampling time of the samples (Figures 5E and 5F). To maximize variability between the reference and query datasets and improve the quality of the reference dataset, we only considered one of the two cell lines in this dataset (Figure 5E). In these analyses, ConDecon inferred a higher abundance for cells that were from the same sampling time than the query bulk sample, even in cases where the bulk sample was from a different patient than the reference single-cell ATAC-seq data (Figures 5F, 5G, and S6D, Pearson’s correlation coefficient between ConDecon’s estimated sampling time and actual sampling time *r* = 0.83, *p*-value = 0.01). The variation in the predicted sampling time was larger during the first 24 hours than in the subsequent 48 hours, suggesting that most chromatin remodeling occurs during the first hours after SOX10 knockdown. However, in these analyses, the performance of ConDecon appeared to be lower than in our studies with RNA-seq data, possibly due to the higher sparsity and presence of batch effects in the single-cell ATAC-seq data.

Altogether, these results show the utility of ConDecon for estimating cell abundances in bulk tissues profiled with other omics data modalities such as spot-based spatial transcriptomics and ATAC-seq data.

### Microglia acquire a *GPNMB*^high^ gene expression phenotype during the mesenchymal transformation of pediatric ependymoma

Having performed a comprehensive evaluation of ConDecon on well-established systems and datasets where the ground truth is known, we next applied ConDecon to a less understood system to assess the potential of ConDecon to discover new biology.

Pediatric ependymoma is a brain cancer that is particularly aggressive in young children due to its frequent relapsing pattern and lack of effective chemotherapies^51-53^. Recent single-cell RNA-seq studies of ependymal tumors have identified a subpopulation of tumor cells with a mesenchymal-like gene expression profile associated with abundant microglia infiltration and poor prognosis^21,22,54,55^. Mesenchymal-like tumor cells in ependymoma are thought to derive from neuroepithelial-like tumor cells by activation of brain injury repair and neuroinflammation pathways in response to microglia-secreted cytokines^21^. To investigate this process, we used ConDecon to study the changes in the gene expression profile of tumor-infiltrating microglia during the mesenchymal transformation of tumor cells. We considered a cohort of 42 ependymal tumors profiled with RNA-seq at the bulk level and a reference single-nucleus RNA-seq atlas of primary and metastatic ependymoma^21^. Our analysis revealed that the abundance of mesenchymal-like tumor cells and microglia in each sample are positively correlated (Pearson’s *r* = 0.70, *p*-value = 3 × 10^−7^), in agreement with previous results^21^. However, it also revealed that tumor-infiltrating microglia experience a continuous transition in their transcriptomic state during the mesenchymal transformation of ependymoma, with different patients showing enrichments of different subpopulations of microglia (Figures 6 and S7). This transition had remained elusive to previous analyses based on conventional gene expression deconvolution approaches^21^. Differentially expressed genes at one end of the microglia trajectory included genes that are characteristic of disease-associated microglia (DAM)^*23-25*^, such as *Apoe, Trem2, Gpnmb, Csf1, Spp1*, and *Il1b*. Using the DAM and mesenchymal gene expression signatures, we introduced a pseudotime in each of the two trajectories describing the transition of neuroepithelial-like tumor cells into mesenchymal-like tumor cells and homeostatic microglia into DAM, respectively (Figure 6A). The ordering of cells along these trajectories was consistent with the RNA velocity vector field^56,57^ (Figure 6B). By computing the expected pseudotime for the microglia and mesenchymal-like tumor cells of each bulk sample based on the probabilities inferred by ConDecon, we found that the state of each sample along the microglia trajectory was strongly correlated with its state along the epithelial-to-mesenchymal-like transition (Figures 6C, 6D, and S7, Pearson’s correlation coefficient between average pseudotime in each trajectory *r* = 0.86, *p*-value < 4 × 10^−13^). Thus, as tumor cells gradually progress from a neuroepithelial-like state onto a mesenchymal-like state, tumor-infiltrating microglia express a DAM gene expression signature consisting of genes involved in phagocytosis and neuroinflammation.

**Figure 6.**
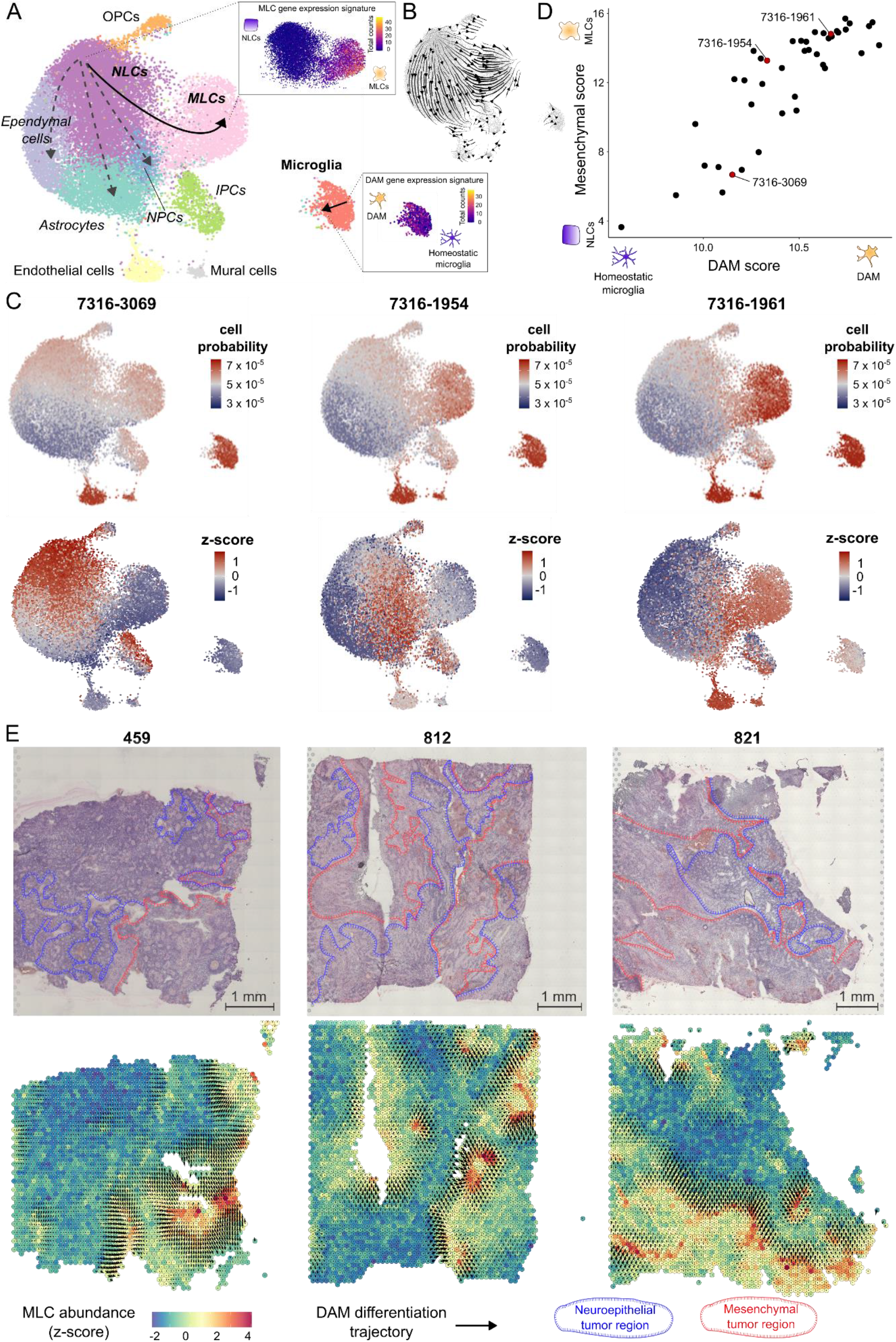
Tumor-infiltrating microglia transition into a disease-associated state during the mesenchymal transformation of pediatric ependymoma. **(A)** UMAP representation of 25,349 cells from 9 posterior fossa ependymal tumors profiled with single-nucleus RNA-seq in Aubin *et al*.^21^. The representation is colored by the annotated cell populations. NLCs: neuroepithelial tumor stem cells; MLCs: mesenchymal tumor cells; NPCs: neural progenitor tumor cells; IPCs: intermediate progenitor tumor cells. The two studied transitions, corresponding to the transformation of NLCs into MLCs and the acquisition of a disease-associated microglia (DAM) phenotype by tumor-infiltrating microglia are schematically indicated. In the inserts, the UMAP representation is colored by the total number of counts of genes belonging to the MLC and DAM gene expression signatures. **(B)** The RNA velocity stream plot showing consistency with the two transitions is shown for reference. **(C)** Single-cell abundance estimates computed with ConDecon for three posterior fossa pediatric ependymal tumors profiled with bulk RNA-seq that span the entire neuroepithelial-to-mesenchymal-like transition. Tumor 7316-3069 has a high abundance of NLCs and small abundance of infiltrating microglia. Most of the microglia are in a homeostatic transcriptional state. In contrast, tumor 7316-1961 has a high abundance of MLCs and infiltrating microglia, and most microglia are in a DAM state. Tumor 7316-490 represents an intermediate state. On the bottom, the same UMAPs are colored by the standardized cell abundances with respect to the full cohort of 42 patients to visualize variability. **(D)** DAM and mesenchymal pseudotimes inferred by ConDecon for 42 posterior fossa pediatric ependymal tumors profiled with RNA-seq at the bulk level. For each tumor, DAM and mesenchymal scores are defined respectively by the average total number of counts of the DAM or MLC gene expression signature inferred by ConDecon for the microglia and tumor cells in each bulk sample. The two scores are correlated (Pearson’s *r* = 0.86, *p*-value < 4 × 10^−13^), indicating that the transition of NLCs into MLCs in the tumor is strongly associated with the transition of infiltrating microglia from a homeostatic transcriptional state onto a DAM state. **(E)** Gene expression deconvolution of three posterior fossa ependymoma tissue sections profiled with spatial transcriptomics. In the bottom, the sections are colored by the relative abundance of mesenchymal tumors cells inferred by ConDecon in each spot. Mesenchymal tumor cells accumulate in localized areas of the tumor. The gradient vector field associated with the microglia DAM pseudotime inferred by ConDecon in each spot is overlaid, showing the transition of microglia into a DAM state in the areas surrounding mesenchymal regions of the tumor. Spatial transcriptomic data from Fu *et al*.^22^. Neuroepithelial and mesenchymal tumor regions have been annotated in the hematoxylin-eosin images according to Fu *et al*.. See also Figure S7.

Mesenchymal tumor cells in pediatric ependymoma localize in perinecrotic regions of the tumor and areas with aberrant vascularization^58^. To characterize the histological organization of the homeostatic-to-DAM transition of tumor-infiltrating microglia in relation to the neuroepithelial-to-mesenchymal transition of ependymoma tumor cells, we used ConDecon to re-analyze a published spatial transcriptomics dataset of pediatric posterior fossa ependymoma^22^. We used ConDecon to infer the mesenchymal tumor cell abundance and microglia pseudotime in each spot of the tissue sections of three tumors using the same reference single-nucleus RNA-seq atlas of primary and metastatic ependymoma^21^. This analysis revealed the accumulation of mesenchymal tumor cells in perinecrotic zones of the tumor and the differentiation of microglia into a DAM state in the regions surrounding them (Figure 6E), adding further support to the inferred relationship between the mesenchymal transformation of ependymoma tumor cells and the homeostatic-to-DAM transition of tumor-infiltrating microglia.

To validate these results, we performed immunohistochemistry on one primary and one metastatic posterior fossa A ependymal tumors. We stained adjacent tissue sections for CA9, which is expressed by mesenchymal-like ependymoma tumor cells^54,59^, IBA1, which is expressed by microglia, and GPNMB, which is expressed by DAM^23^. Consistent with the predictions of ConDecon, the immunohistochemistry data showed that microglia surrounding or infiltrating mesenchymal regions of the tumors expressed high levels of the DAM marker GPNMB (Figures 5E and S6). In contrast, we did not detect the expression of GPNMB in microglia infiltrating non-mesenchymal regions of the tumors (Figure 7).

**Figure 7.**
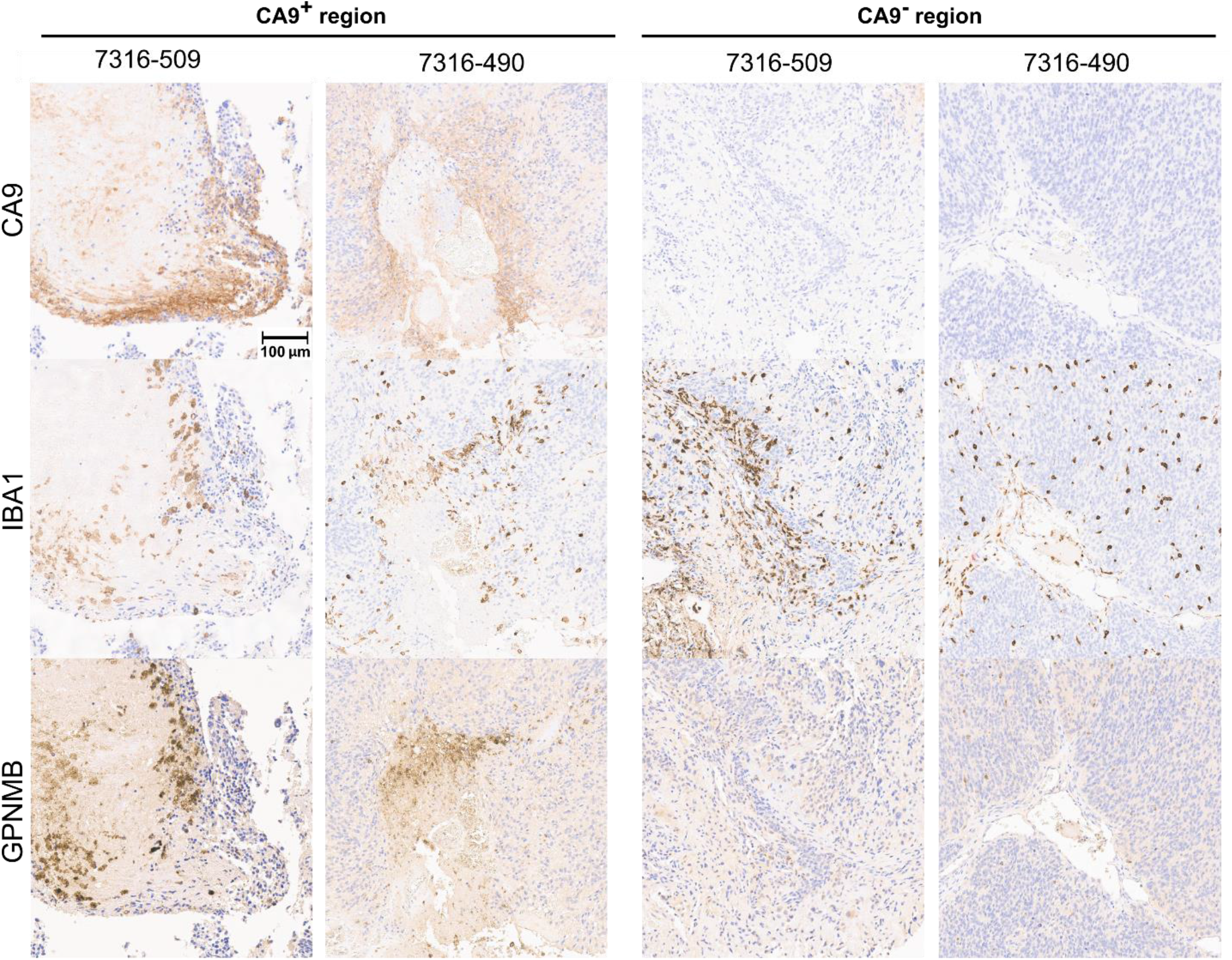
Immunohistochemistry of infiltrating microglia in pediatric ependymal tumors. Immunohistochemistry staining of adjacent tissue sections from two pediatric ependymal tumors (7316-509 and 7316-490). Each tumor was stained for CA9 (a marker of mesenchymal regions), IBA1 (a microglial marker), and GPNMB (a DAM marker). Consistent with the predictions of ConDecon (Figure 6), microglia surrounding CA9^+^ mesenchymal regions of the tumor acquired a DAM state, marked by the expression of GPNMB, while microglia in neuroepithelial regions of the tumor do not have detectable levels of GPNMB. Scale bar: 100 μm.

In summary, these findings indicate that key microglia inflammatory pathways play a role in the mesenchymal transformation of pediatric ependymoma. Of note, a recent work has shown evidence for the involvement of GPNMB-high microglia/macrophages in the mesenchymal transformation of glioblastoma^60^, suggesting that GPNMB-high microglia/macrophages may play a role in the mesenchymal transformation of multiple gliomas. Altogether, these findings showcase the capability of ConDecon to uncover new biological insights by using various transcriptomic datasets.

## Discussion

Estimating cell abundances in bulk tissues has been critical to addressing questions related to cellular heterogeneity using bulk transcriptomic data. Although current methods for gene expression deconvolution provide robust and accurate estimates of cell abundances for discrete cell types, they are limited in their ability to infer changes derived from continuous and dynamic cellular processes such as cell differentiation, immune cell activation, or wound healing. The emergence of single-cell RNA-seq technologies in the past decade has provided new powerful avenues for studying questions of cellular heterogeneity in tissues. However, the scalability and applicability of single-cell RNA-seq remains limited. Here, we presented ConDecon, a conceptually different approach to gene expression deconvolution that can detect fine-resolution changes in cell abundance from bulk tissues using single-cell RNA-seq data as a reference.

ConDecon conceives the bulk tissue as being generated by a stochastic sampling process where cells from the reference single-cell dataset are sampled with different probabilities, and it infers the probability for each cell in the reference dataset to be present in the bulk tissue. The approach thus requires the reference single-cell dataset to be representative of the cell states that are present in the bulk tissue but not necessarily of their cell abundances.

Our analyses using both actual and simulated data demonstrate that, like current methods for gene expression deconvolution, ConDecon can accurately estimate cell abundances associated with discrete cell types. However, in contrast to those methods, it can also recapitulate gradual changes in cell state that would otherwise be obscured by conventional clustering-based approaches. We have demonstrated the potential of this type of inference in biomedical applications by reanalyzing published data of pediatric ependymal tumors, where we have discovered the implication of microglial neurodegenerative programs in the mesenchymal transformation of these tumors. These results indicate the involvement of DAMs in the mesenchymal progression of pediatric ependymoma. Furthermore, they demonstrate the potential of ConDecon to reveal new biological insights by utilizing diverse transcriptomic datasets. In this regard, we have also shown the adaptability of ConDecon’s approach to other omics data modalities, like chromatin accessibility data, for which there is currently a scarcity of deconvolution approaches. We anticipate that these features will improve our understanding of cellular heterogeneity and tissue cell composition by greatly facilitating the inference of cell state abundances within complex bulk tissues, particularly in the context of evolving systems like development and disease progression.

## Limitations

Throughout our benchmarking analyses we have discussed several limitations of ConDecon. Most importantly, ConDecon requires the user to provide a reference single-cell RNA-seq dataset of the same tissue type as the query bulk tissue. Our analyses using real data with known ground-truth cell type abundances (Figure S3) show that ConDecon cannot infer very large departures from the populations and abundances present in the reference single-cell RNA-seq dataset. Using a reference dataset that largely differs from the query bulk data will therefore lead to incorrect estimates, as demonstrated in Figure S3D. However, given the current broad availability of single-cell data from most tissues, we do not expect this to be a major limitation in practical cases.

Relatedly, in some datasets, the inferred cell type abundances by ConDecon have a limited range compared to the ground-truth abundances (Figure 2A). Nonetheless, in these cases, we observe that the inferred and ground-truth abundances are still strongly correlated, enabling the inference of statistical associations with other variables, such as clinical variables.

Lastly, the simulation of bulk RNA-seq data from single-cell RNA-seq data is another potential venue for improving ConDecon. Our current approach is based on aggregating single-cell RNA-seq counts across cells to build pseudo-bulk datasets. Although this approach is standard in the field and produces good results, it does not explicitly model the technical differences between single-cell and bulk RNA-seq technologies. Developing more realistic simulations of bulk RNA-seq data based on single-cell RNA-seq data will improve the estimates of ConDecon.

We expect that continued work in these directions will result in improved versions of ConDecon in the coming years.

## Supporting information

Key Resources

## Acknowledgments

The authors are grateful to the CBTN for providing data and tissue specimens for conducting this study and to the Pathology Core of the Children’s Hospital of Philadelphia, Dr. Alice-Chen-Plotkin, Eliza Brody, and Marc Carceles-Cordon for excellent technical assistance with the immunohistochemistry staining of FFPE tissue sections. The work of P. G. C. and R. G. A. has been partially supported by the Pediatrics Networks for the Human Cell Atlas from the Chan Zuckerberg Initiative. The work of P. G. C. has also been partially supported by the U. S. National Institutes of Health (NIH) through the grant U01CA269409-01A1 from the National Cancer Institute (NCI) Informatics Technologies for Cancer Research (ITCR) program. The work of J. M. has been supported by the NIH grant R25HG010323-01A1 from the National Human Research Institute (NHGRI).

## Author contributions

R. G. A. implemented ConDecon. R. G. A., J. M., and R. H. conducted the computational analyses. P. G. C., E. G., and P. N. formalized ConDecon mathematically. R. G. A. and P. G. C. conceived the project and wrote the first draft of the manuscript.

## STAR Methods

### Resource availability

#### Lead contact

Further information and requests for resources and reagents should be directed to and will be fulfilled by the lead contact, Pablo G. Camara.

#### Materials availability

This study did not generate new unique reagents.

#### Data and code availability

- This paper analyzes existing, publicly available data. The accession numbers for the datasets are listed in the key resources table.
- Full, unedited IHC images have been deposited in Mendeley Data. Accession numbers are listed in the key resources table.
- All original code has been deposited at GitHub and is publicly available as of the date of publication. DOIs are listed in the key resources table.
- Any additional information required to reanalyze the data reported in this paper is available from the lead author upon request.

### Method details

#### Overview of ConDecon

ConDecon uses the count matrix and latent space of the reference single-cell RNA-seq dataset to estimate the likelihood for each cell in the dataset to be present in the query bulk tissue sample. For that purpose, it considers the set 𝒢 of genes included in both the single-cell and bulk gene expression datasets and the subset *𝒯* ⊂ 𝒢 of most variable genes used to build the single-cell gene expression latent space. For each cell in the single-cell RNA-seq dataset, ConDecon aggregates the gene expression counts of the cell and its *r* nearest neighbors in the latent space (default *r =* 5) and computes the Spearman’s correlation between the bulk and the aggregated cells among the *𝒯* genes, where genes with tied expression values are assigned their average rank. We denote by 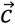 the vector of correlation coefficients computed in this manner across all cells in the single-cell dataset.

The goal of ConDecon is to infer a vector 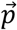 of cell probabilities starting from 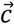. For that purpose, it is convenient to expand 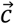 and 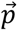 in an orthonormal basis of functions with support on the latent space of the single cell dataset,

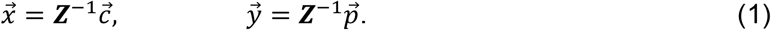

In these expressions, ***Z*** is a *J* by *D* matrix containing the cell loadings in the latent space, ***Z***^−1^ is its Moore-Penrose inverse, and 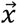 and 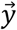 are vectors in *D* dimensions, where *J* is the number of cells in the single-cell dataset and *D* is the number of dimensions of the latent space (default *D =* 10). By reducing the dimensionality of the problem in this manner, we facilitate learning the relationship between 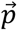 and 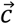,

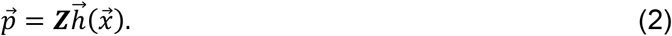

In our analyses, we use the Seurat pipeline^61^ to compute ***Z***, where the single-cell gene expression matrix is log-normalized by library size, restricted to the top *T*most variable genes (default *T=* 2,000), and reduced to *D* dimensions by Principal Component Analysis (PCA).

ConDecon uses a polynomial model of degree *s* for 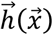 (default *s =* 1),

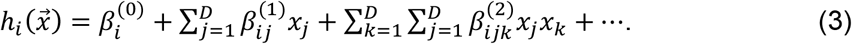

To estimate the coefficients *β*, it generates a training dataset consisting of *k* simulated cell abundance distributions in the latent space (default *k =* 5,000),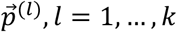. Any family of probability distributions that can approximate any given smooth probability distribution with support on the latent space with arbitrary precision suffices for modeling the cell abundance distributions. ConDecon uses a mixture of *M*^(*l*)^ *D*-dimensional Gaussian distributions due to its easy implementation,

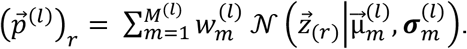

where 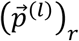 denotes the *r*-th component of the *J*-dimensional vector 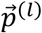, corresponding to the probability of cell *r* in the simulated dataset *l*, and 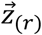 is the *D*-dimensional vector given by the *r*-th row of ***Z***, corresponding to the coordinates of the cell *r* in the latent space. The number of components, *M*^(*l*)^, is uniformly sampled from a range of values (default *M*^(*l*)^ ∈ [1,5]); the location of each center 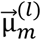 in the latent space is given by a randomly sampled cell from the single-cell data; the covariant matrix 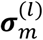 is taken to be proportional to the identity with proportionality constant uniformly sampled from a finite range of values such that the fraction of cells within two standard deviations of the center is in a given percentile range (default 5-20%); and each mixing parameter 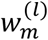 is uniformly sampled from [0,1]. The training data thus consists of a rich set of probability distributions containing a varying number of components spanning diverse locations across the entire latent space. For each probability distribution 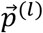, *n* cells are sampled (with replacement) from the single-cell data (default 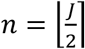), their gene expression counts are aggregated to create a synthetic bulk gene expression profile, and a vector of correlation coefficients 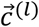 is computed as described above. The pairs 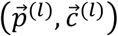 are then transformed into pairs 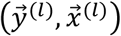 using equations (1), and they are used to estimate the coefficients *β* in equation (3) using linear least squares regression.

Since ConDecon is trained on probability distributions, most of the values in the vector 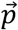 inferred by ConDecon (equation (2)) are between 0 and 1. However, since the map in equation (3) is unconstrained, there may be cells with close to zero but negative probabilities in some situations. We address these cases by normalizing the final vector of probabilities as,

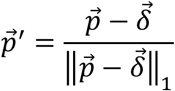

where 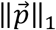 denotes the *L*^1^-norm of 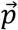 and 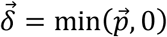 is a vector with all entries equal to the smallest negative element in 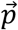 or 0 if all elements are positive. This normalization ensures the output probabilities are between 0 and 1 and add up to 1. We empirically find that this approach offers slightly better results in practical situations than setting negative values to zero.

#### Theoretical foundation of ConDecon

We now discuss the theoretical justification of several aspects of the approach implemented in ConDecon:

##### Simulated vs real bulk transcriptomic data

A key aspect in deconvolving bulk RNA-seq data using reference single-cell RNA-seq data is the ability to compare the gene expression profiles produced with these two technologies. The standard approach aggregates the gene expression counts of the cell populations of interest in the single-cell data to produce “synthetic” bulk transcriptomic data that can be used to deconvolve the actual bulk dataset^10,12,35,62^. However, the gene expression values in the synthetic bulk datasets still can differ substantially from the expression values that would result from profiling the same sample of cells with actual bulk RNA-seq due to the large technical differences between bulk and single-cell RNA-seq, therefore limiting the accuracy of the deconvolution. To be less sensitive to those technical differences, ConDecon uses gene rank correlations instead of gene expression levels to estimate cell abundances. To verify that the gene ranks can better discriminate biological differences between samples than gene expression values in synthetic bulk data, we considered two datasets consisting of paired single-cell or single-nuclei and bulk RNA-seq data from the same samples, encompassing bone marrow samples from 8 patients^28^, tumor samples from 8 high-grade serous ovarian cancer patients^63^, and tumor samples from 17 Wilms tumor patients from the scPCA. For each sample, we aggregated the single-cell or single-nuclei counts across all the cells to construct a synthetic bulk RNA-seq dataset for the sample. We then used Pearson correlation across the top 2,000 variable genes to compare the expression levels or the gene ranks between the synthetic and the actual bulk RNA-seq datasets (Figure S1A). This analysis revealed a higher correlation between the ranks of the genes than between the expression values of the genes constructed in this manner (Figure S1A). Furthermore, the correlation between gene ranks in synthetic and actual bulk RNA-seq data from the same patient was significantly higher than the correlation observed between synthetic and actual bulk data from different patients (Wilcoxon rank-sum test *p*-value < 0.05 in the three datasets). These results show that the gene ranks obtained from aggregating single-cell or single-nuclei counts can discriminate samples from the same tissue type but from different patients. In contrast, the correlation between gene expression values in synthetic and actual bulk RNA-seq data from the same patient was not significantly higher than the correlation between synthetic and bulk data from different patients in two of the three datasets (Figure S1A), indicating limited power to distinguish samples from the same tissue type but different patients based on aggregated single-cell gene expression levels.

##### Uniqueness of solutions

A potential limitation of using rank correlation to infer cell abundances from bulk gene expression profiles is that different cell abundance configurations can lead to the same vector of correlation coefficients 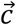. This means that 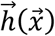 is multivalued, even without collinearity in the reference single-cell gene expression matrix. However, this concern can be safely disregarded when working with single-cell datasets that consist of hundreds of variable genes. To understand why, consider the expression table *G* of a reference single-cell dataset consisting of *J* cells and *T*variable genes. A vector of cell abundances 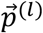 represents a point in a regular (*J*-1)-simplex Ω, with volume given by the formula 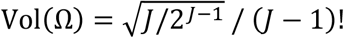. The gene expression profile of the synthetic bulk dataset corresponding to 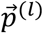 is given by the vector 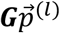, which induces an ordering on the variable genes. Vectors of cell abundances that lead to the same ordering of genes cannot be differentiated by ConDecon. Thus, we can think of Ω as partitioned into a finite number of contiguous subspaces or “tiles”. The number of tiles is given by the number of distinct gene orderings induced by 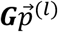 as 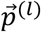 varies across Ω. This number is at most *T*!. Consequently, in real-world scenarios, the volume of each tile in Ω is very small, and the accuracy of cell abundance inferences is limited by other factors rather than the partition of Ω.

The space of all possible gene orderings can be represented by the set of all permutations of the elements in the tuple (1, 2, …, *T*). In mathematical terms, this corresponds to the symmetric group of degree *T, S*_*T*_. We can endow this space with a metric by considering the normalized Kendall’s *τ*rank distance^64,65^,

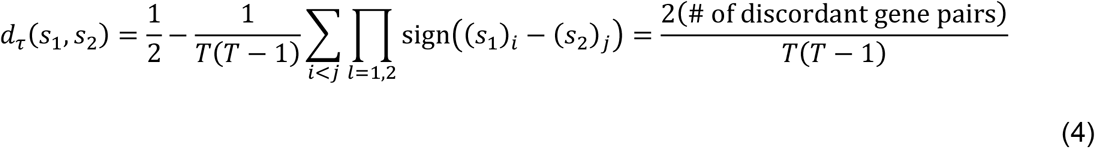

where *s*_1_, *s*_2_ ∈ *S*_*T*_, sign(*x*) denotes the sign of *x*, and discordant gene pairs refer to variable genes with different relative ordering between *s*_1_ and *s*_2_. In particular, *d*_*τ*_(*s*_1_, *s*_2_) satisfies all the axioms of a distance function, including the triangle inequality^**64**,**65**^.

The distance in *S*_*T*_between the synthetic bulk gene expression profiles induced by two vectors of cell abundances, 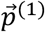 and 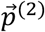, is thus given by 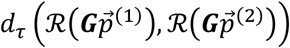, where 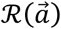 gives the element of *S*_*T*_corresponding to the gene ranks in the gene expression vector 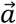 according to a predefined ordering operation (e.g., smaller to greater). When the number of variable genes *T*is large, we empirically observe that the expected value and variance of 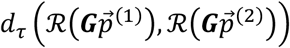 for fixed *G* and uniform sampling from Ω are approximately proportional to the root mean square error (RMSE) between 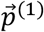 and 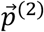 (Figures S1B and S1C). In this scenario, 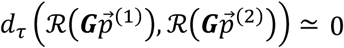 if and only if RMSE 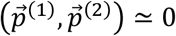 and any configurations of cell abundances leading to the same vector 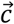 of correlation coefficients are similar to each other. Propositions 1 and 2 below offer a theoretical justification for this observation.

##### Mathematical derivation

We now present the theoretical basis of ConDecon in precise mathematical terms. Let {*G*_*k*_; *k =* 1, …, *J*} be a set of independent absolutely continuous random variables, not necessarily identically distributed, with probability density functions ρ_*k*_, and let {*G*_*ik*_ ∈ ℝ; *i =* 1, …, *T*} be a set of *T*values sampled from *G*_*k*_. For convenience, we arrange the values *G*_*ik*_ into a *T*× *J* matrix *G* (the single-cell gene expression table). Without loss of generality, we assume that rank(*Ĝ) = T*, where *Ĝ* is the matrix that results from applying ℛ to the columns of *G*. Note that given some matrix *G*, it is always possible to construct a smaller matrix by Gaussian elimination that satisfies the condition.

Let 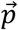 be a point in the (*J* − 1)-dimensional probability simplex 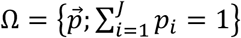 (the space of cell abundance distributions). We define the following map,

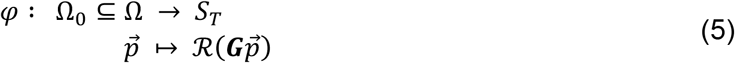

where Ω_0_ ⊆ Ω is the subspace for which 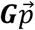 has no ties in the ranking of its elements. By construction, the map φ induces an injection from the set of open sets {ω_*s*_} into *S*_*T*_, where the open sets are given by,

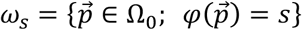

where Ω_0_ *=* ⊍_*s*_ ω_*s*_ and ⊍ denotes the disjoint union. From equation (5), we observe that,

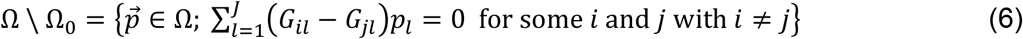

where *i* and *j* can take values 1, …, *T*. In particular, Ω \ Ω_0_ has zero measure in Ω with probability 1.

The following proposition forms the conceptual basis of ConDecon:

###### Proposition 1

For any pair of points 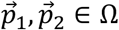, with probability 1 there is a sufficiently large *T*for which they are separated by Ω \ Ω_0_.

*Proof*. Let 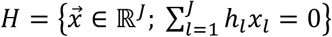 be a hyperplane that separates 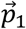 and 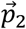 and choose a sufficiently small ε > 0 such that any hyperplane 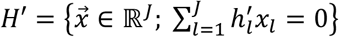 with 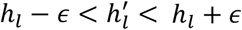 also separates 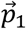 and 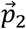. From equation (6), the problem is then reduced to finding *G*_*il*_ and *G*_*jl*_ such that *h*_*l*_ − ε < *G*_*il*_ − *G*_*jl*_ < *h*_*l*_ + ε.

Let ***A***_*l*_ denote the random variable defined by *G*_*l*_ − *G*_*l*_, where the two terms in the subtraction correspond to independent and identically distributed copies of the random variable *G*_*l*_. Using the convolution of probability distributions, we note that the probability density function of **A**_*l*_ is given by

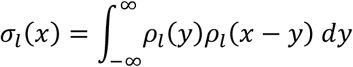

In particular, σ_*l*_(*x*) *=* 0 if and only if ρ_*l*_(|*x*|*) =* 0. Hence,

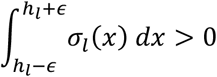

where, without loss of generality, we assume that all the *h*_*l*_ and ε have been rescaled by the same factor so that (*h*_*l*_ − ε, *h*_*l*_ + ε) ⊆ *supp(***A**_*l*_). Therefore, with sufficient sampling we can always find *G*_*il*_ and *G*_*jl*_ such that *h*_*l*_ − ε < *G*_*il*_ − *G*_*jl*_ < *h*_*l*_ + ε. ◼

From equation (5), we then conclude that (*S*_*T*_, *d*_*τ*_) can be isometrically embedded into Ω.

To invert the map φ using machine learning approaches, it is convenient to work with a Euclidean embedding of *S*_*T*_. This embedding can be constructed with the use of distance coordinates^66^,

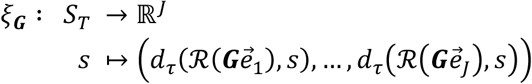

where 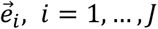, are *T*-dimensional vectors with all entries equal to zero except for the *i*-th element, which is equal to 1. We refer to this embedding as the space of gene rank correlations. The ConDecon injection is then defined as *h*^−1^ *≡* ξ_*G*_ ∘ φ.

In practical situations, we find it convenient to approximate Kendall’s *τ*rank distance using the Spearman rank correlation coefficient *ρ*, where 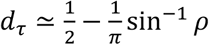 for a normal population, as ρ can be computed in *O(Tlog T*) time instead of *O(T*^2^).^67^

##### Regularization

Collinearity in the reference gene expression data poses an inherent limitation to gene expression deconvolution methods. Regression-based methods produce degenerate solutions when the reference gene expression signature matrix exhibits substantial collinearity, necessitating regularization schemes. In ConDecon, collinearity in the reference single-cell gene expression matrix *G* results in a reduction in the number of gene orderings induced by 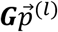. This decrease leads to an increase in the volume of the tiles in Ω, which leads to an increase in the uncertainty associated with the estimation of cell abundances. In terms of 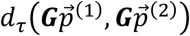, collinearity reduces the magnitude of the proportionality constant for the expected value of 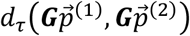 as a function of RMSE 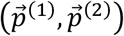 without substantially altering the variance (Figures S1B and S1C). However, when the number of variable genes is large, 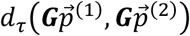 is relatively robust against the presence of collinearity. For instance, by artificially increasing the amount of collinearity in published single-cell datasets by replacing the gene expression profile of some of the cells with rescaled copies of the expression profile of other cells in the dataset, 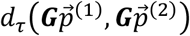 continues to be small if and only if RMSE 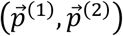 is small, even when the rank of *G* is reduced by two orders of magnitude using this procedure (Figures S1B and S1C).

If the gene expression profiles of a set of cells *𝒞*is approximately collinear, configurations of cell abundances 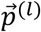 that only differ in the abundance of cells in *𝒞*will belong to the same tile of Ω, effectively reducing the number of tiles in Ω. Since the cells *𝒞*are in the same region of the single-cell gene expression latent space, it is possible to regularize the inference of cell abundances by expanding the cell abundances 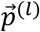 into principal or diffusion components of the latent space and keeping only the first *D ≪ J* terms in the expansion,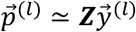. By only considering configurations of cell abundances that vary smoothly on the latent space, the effective dimensionality of Ω is reduced without substantially changing the number of tiles, and the stability of 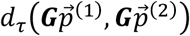 against collinearity is improved as a function of RMSE 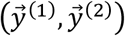 (Figure S1B).

#### Deconvolution of simulated gene expression data

We used the R package splatter^27^ (v1.10.1) to simulate single-cell RNA-seq data with either a distinct number of cell types (splatSimulateGroups) or a continuous cell differentiation trajectory (splatSimulatePaths). Each simulation contained 5,000 cells (batchCells), 20,000 genes (nGenes), approximately 45 positively differentially expressed genes per group, and approximately 5 negatively differentially expressed genes per group (de.prob = 0.0025, de.downprob = 0.1).

##### Simulation of discrete cell types

To simulate discrete cell types, we generated synthetic single-cell data containing either 3, 6, or 9 cell types of equal size (group.prob) and 9 levels of differentially expressed genes (de.facLoc ∈ [0.01, 0.05, 0.1, 0.15, 0.2, 0.3, 0.4, 0.5, 0.6]). For each of the 27 simulated single-cell datasets, we generated 25 corresponding bulk gene expression profiles by aggregating cells from each cell type *k* with varying proportions 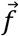. For that purpose, *n*_*k*_ cells were uniformly sampled (without replacement) from each cell type *k*, such that 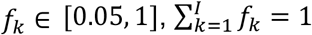 and 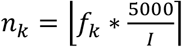 where *f*_*k*_ is the simulated abundance of cell type *k* and *I* is the total number of cell types. For each of the 27 simulations, we ran ConDecon with default parameters using the top 10 principal components and 2,000 variable genes calculated with the R package scran^68^ (v1.14.6). We aggregated ConDecon’s inferred cell probabilities 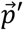 into inferred cell type abundances 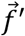,

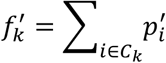

and compared them with the simulated cell type abundances *f*_*k*_ by computing their Pearson correlation and RMSE across samples (cell-type-level performance) or cell types (sample-level performance).

##### Simulation of cell differentiation trajectories

We simulated single-cell data of cell differentiation trajectories with three different topologies (a tree with a bifurcation (path.from = c(0, 1, 1)), a tree with a three-way split (path.from = c(0, 1, 1, 1)), and a tree with two consecutive bifurcations (path.from = c(0, 1, 1, 3, 3))) and 13 levels of differential gene expression (de.facLoc ∈ [0.01, 0.05, 0.1, 0.15, 0.2, 0.3, 0.4, 0.5, 0.6, 0.7, 0.8, 0.9, 1], such that each branch of the trajectory is approximately straight (path.sigmaFac = 0.5), has genes expressed in a nonlinear manner along the path (path.nonlinearProb = 0.3), and is composed of approximately the same number of cells (group. prob). Pseudotime was interpolated across each trajectory such that there were approximately 20 cells in each iterative step of pseudotime (path.nSteps). For each of the 39 simulated single-cell datasets, we generated 25 bulk gene expression profiles by aggregating 1,000 cells sampled from the single-cell data (with replacement) based on a randomly generated Gaussian distribution *N*(μ, σ) where µ is a uniformly sampled cell along pseudotime and σ is uniformly sampled from a range of sigma values that are calculated to on average capture 300 - 1,500 cells within two standard deviations of a center. For each of the 39 simulations, we ran ConDecon with default parameters using the top 10 principal components and 2,000 most variable genes calculated with scran (v1.14.6). To evaluate performance, we calculated the Pearson’s correlation coefficient and RMSE between ConDecon’s inferred cell probabilities and the simulated ground truth cell abundances.

#### Comparison to clustering-based methods for gene expression deconvolution

We used the benchmarking pipeline of Avila-Cobos *et al*.^32^ to evaluate the ability of ConDecon and 17 other deconvolution methods to infer discrete cell type abundances in bulk tissues. This pipeline builds synthetic bulk RNA-seq datasets by aggregating the gene expression counts of cells sampled from real single-cell RNA-seq datasets. We used 6 single-cell RNA-seq datasets^10,11,28,33,34,69^, which we filtered using the same quality control steps outlined in Avila-Cobos *et al*.^32^. In brief, we filtered out genes expressed in less than 5% of the cells, and cells with a total, mitochondrial, or ribosomal UMI count greater than 3 deviations from the median across all the genes. We only considered cell types with at least 50 cells. We down sampled the bone marrow dataset to 8,000 cells. Datasets were then processed using the Seurat pipeline^61^. Cells were log-normalized by library size and the top 2,000 most variable genes were selected for PCA. For datasets that contained more than one sample (all except for Newman *et al*.^10^), we used Harmony with default parameters to consolidate the top 30 principal components across samples. Cells were clustered using Louvain community detection based on the top 30 latent dimensions and cell populations were annotated using the same sets of markers as in the original papers. As described in Avila-Cobos *et al*.^32^, each single-cell dataset was then subset into two equal-sized datasets representing a reference single-cell dataset and a single-cell dataset that was used to build 1,000 synthetic bulk query datasets. Each synthetic bulk query dataset was constructed by aggregating the gene expression counts of *n* cells sampled from each cell type with different proportions, where the number of sampled cells depended on the size of the dataset (Oetjen *et al*.^28^, *n =* 4,000; Baron *et al*.^11^, *n =* 3,500; Enge *et al*.^33^, *n =* 1,000; Han *et al*.^34^, *n =* 3,000; Newman *et al*.^10^, *n =* 3,500). For each dataset, either none, a small, a medium, or a large cell type was removed from the reference single-cell data to evaluate the stability of the estimates against missing data (Oetjen *et al*.^28^: non-classical monocytes (small, 1.6%), CD14 monocytes (medium, 8.0%), T cells (large, 42.8%); Baron *et al*.^11^: delta cells (small, 7.0%), ductal cells (medium, 12.6%), beta cells (large, 29.5%); Enge *et al*.^33^: beta cells (small, 15.0%), acinar cells (medium, 17.6%), alpha cells (large, 44.7%); Han *et al*.^34^, loop of Henle thick ascending limb cells (small, 10.7%), proximal tubule cells MT1G high (medium, 18.3%), intercalated cells (large, 28.7%); Newman *et al*.^10^: NK T cells (small, 4.4%), CD8 T cells (medium, 21.5%), CD4 T cells (large, 27.2%). We quantified the performance of each algorithm across each dataset and condition by computing the average Pearson’s correlation coefficient, average Lin’s correlation coefficient, and root mean squared error of the predicted values in comparison with the simulated ground-truth values, combined across all samples and cell types. In these analyses, we ran ConDecon with default parameters using the top 10 latent dimensions and 2,000 most variable genes computed with Seurat (v3.1.5). We aggregated ConDecon’s inferred cell probabilities into inferred cell type proportions as described above. All the other gene expression deconvolution methods were run using the marker genes and default parameters specified in the original publication of the benchmarking pipeline^32^ and the associated code repository (https://github.com/favilaco/deconv_benchmark).

#### Comparison of estimated cell type abundances to FACS data

We benchmarked ConDecon, CIBERSORTx^10^ (S-mode), MuSiC^12^, Bisque^35^, and CPM^37^ using two bulk RNA-seq datasets of human bone marrow^28^ and PBMC^10^ for which paired FACS data were available. In these analyses, ConDecon and all the other algorithms were run with default parameters. CIBERSORTx, MuSiC, and Bisque were used with the marker genes output by the “Create Signature Matrix” tool of CIBERSORTx. CPM was run using quantifyTypes = TRUE and a homogeneous 2-dimensional representation, as recommended in its documentation (https://github.com/amitfrish/scBio). To build the homogeneous 2-dimensional representation for CPM, we applied UMAP to the top 30 principal components based on the top 2,000 least variable genes expressed in > 5% of cells. For each bulk sample, the cell weights inferred by CPM were shifted by their minimum value so that they were all positive, and then normalized as probabilities and aggregated into cell type proportions. To evaluate the performance of each algorithm, we calculated the Pearson’s correlation coefficient, Lin’s correlation coefficient, and RMSE between the predicted cell type proportions and the FACS data across bulk samples (cell-type-level performance) or cell types (sample-level performance).

#### Comparison of estimated cell type abundances to snRNA-seq data

We benchmarked ConDecon, CIBERSORTx^10^ (S-mode), MuSiC^12^, Bisque^35^, and CPM^37^ using two bulk RNA-seq datasets of human Wilms tumors and pediatric glioma from the Alex’s Lemonade Stand Single-Cell Pediatric Cancer Atlas (scPCA) for which paired single-nucleus RNA-seq data were available (Wilms tumors, *n* = 17; pediatric glioma, *n* = 37). Since single-nucleus RNA-seq is not biased by the size and shape of cells during droplet encapsulation, we reasoned that these data would provide a good estimation of cell type proportions for the bulk RNA-seq data in situations where FACS data is not available due to the absence of well-validated antibody panels specific to the tissue.

We downloaded the filtered gene expression data from the scPCA portal. Ensembl IDs were converted to Hugo gene names using the biomaRt database (version 2.50.3). For the single-nucleus RNA-seq data of Wilms tumors, we excluded two samples (SCPCL000003 and SCPCL000018) with a median number of expressed genes per cell < 300. We filtered out cells with < 500 expressed genes, < 750 UMIs, > 75,000 UMIs, or > 25% UMIs corresponding to mitochondrially encoded genes. For the single-nucleus RNA-seq data of pediatric central nervous system tumors, we excluded one sample (SCPCL000545) with less than 100 cells. We filtered out non-demultiplexed cells (labeled as NA in the original dataset), as well as cells with < 500 expressed genes, < 750 UMIs, > 50,000 UMIs, or > 10% UMIs corresponding to mitochondrially encoded genes. We down sampled the pediatric central nervous system tumor dataset to 10,000 cells. Single-nucleus RNA-seq datasets were then processed using the Seurat pipeline^61^. In particular, cells were log-normalized by library size and the top 2,000 most variable genes were selected for PCA. We used Harmony with default parameters to consolidate the top 20 principal components across the samples. Cells were clustered using Louvain community detection based on the top 20 latent dimensions. To benchmark ConDecon, CIBERSORTx, MuSiC, Bisque, and CPM, we used the same procedure outlined in subsection “Comparison of estimated cell type abundances to FACS data”, with ground truth cell proportions derived from the single-nucleus RNA-seq data cell type proportions for each sample.

#### Analysis of the stability of cell abundance estimates using real RNA-seq data

We varied ConDecon’s input parameters through a broad range of values in the deconvolution of the human bone marrow^28^ and PBMC^10^ datasets. In this analysis, we systematically evaluated the Pearson’s correlation and RMSE between the estimated cell type proportions and FACS data across samples (cell-type-level performance) or cell types (sample-level performance) while varying one parameter and keeping the remaining parameters steady. We varied the number of variable genes *T*(100, 300, 500, 1,000, 2,000, 4,000) used to compute gene ranks while keeping the number of principal components *D =* 10 and the degree of the polynomial *s =* 1; the number of principal components *D* (2, 4, 6, 8, 16) while keeping *T=* 2,000 and *s =* 1; and the degree of the polynomial *s* (1, 2) while keeping *T=* 2,000 and *D =* 10. In addition, we down-sampled the number of cells in the reference single-cell datasets uniformly (10%, 25%, 100%) and non-uniformly (to mimic the cell type proportions in the reference single-cell data, the average cell type proportions in the FACS data, and equal cell type proportions) while keeping *T=* 2,000, *D =* 10, and *s =* 1. For the non-uniform sampling, the bone marrow and PBMC datasets were restricted to 700 and 1,630 cells, respectively. These values were determined by the number of cells in the smallest cell population in the single-cell RNA-seq data for which FACS data was available, so that we could generate datasets with equal cell type proportions without having to remove any cell types.

We also used the two datasets to evaluate the stability of ConDecon’s cell abundance estimates across 20 random seeds. For each cell type, we calculated the standard deviation of the cell type abundance estimates across the 20 initializations for each query bulk RNA-seq sample and compared it with the standard deviation of the cell type abundance estimates across query bulk RNA-seq samples for each initialization.

Finally, we considered single-cell RNA-seq data of the mouse kidney (*n =* 16) and bone marrow (*n =* 13) from the Tabula Muris Senis atlas^29^. The gene expression count tables from the two tissues were concatenated together and processed using the Seurat pipeline^61^. Cells were log-normalized by library size and the top 5,000 most variable genes were selected for Principal Component Analysis (PCA). Cells were clustered using Louvain community detection based on the top 20 principal components. We then considered 54 mouse bone marrow samples profiled by bulk RNA-seq from the Tabula Muris Senis atlas^29^. We removed one bulk sample that had < 1,000 expressed genes from downstream analyses. We then used the concatenated single-cell RNA-seq data as reference for ConDecon to deconvolve the mouse bone marrow bulk RNA-seq data. We ran ConDecon using the top 20 principal components and 5,000 most variable genes. To understand the importance of using a representative reference dataset when running ConDecon, we deconvolved the bone marrow bulk RNA-seq data using single-cell RNA-seq data from either the bone marrow dataset alone or from the kidney and bone marrow datasets concatenated together. We reasoned that running ConDecon using reference data derived from the same tissue type that the query bulk dataset would result in training data that better resembles the query data, as defined by their separation in the space of probability distributions *Y*. To test this, we defined a proximity score for each query bulk sample using both reference datasets. The proximity score is calculated by taking the Euclidean distance between each query sample and the 10 nearest training points in *Y*, normalized by the average Euclidean distance between the 10 nearest training points in *Y*.

#### Analysis of B cell maturation

We considered the longitudinal mouse bone marrow single-cell RNA-seq dataset from the Tabula Muris Consortium^29^, which consists of 13 and 53 mice profiled at the single-cell and bulk level, respectively. We followed the same quality control procedures described above (subheading “Comparison to clustering-based methods for gene expression deconvolution”) to process the single-cell RNA-seq data, with the addition of removing genes expressed in < 0.1% of the cells. We used Seurat^61^ (v3.1.5) to log-normalize the gene expression profile of each cell by library size and perform PCA using the top 2,000 variable genes. We then consolidated the single-cell data of the 13 mice using Harmony with default parameters and the top 30 principal components. The resulting consolidated latent space was clustered using Louvain community detection and the clusters were annotated using the marker genes identified in the original reference. The cell population annotated as “hematopoietic stem cells” in the original publication was reannotated as hematopoietic stem and progenitor cells (HSPCs) based on the expression of hematopoietic progenitor markers such as *Mpl, Ctsg*, and *Gata1*^70^. We used Monocle 3^71^ (v0.2.3) with default parameters to infer a cell differentiation pseudotime 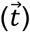 in the single-cell gene expression space of the B cell lineage. We then ran ConDecon, CIBERSORTx (S-mode), MuSiC, Bisque, and CPM as described above (subheading “Comparison of estimated cell type abundances to FACS data”). We evaluated the performance of these methods by calculating the log_2_-fold change of the median predicted cell type proportion of each B cell subpopulation between samples from young (≤ 3 months) and adult (> 3 months) mice. Statistical significance was calculated using a Wilcoxon rank sum test. In addition, we used the single-cell probabilities 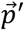 inferred by ConDecon to compute the estimated average pseudotime of the B cells in each bulk sample 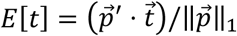, where only the interquartile range of the distribution of probabilities was considered in the estimation. We then tested the association between the estimated average pseudotime and the mouse age of each sample using Pearson’s correlation. We repeated the same analysis using MeDuSA^41^ v1.0, with parameter `fractional = TRUE`, to estimate the relative cell abundances along the B-cell lineage for each bulk sample.

#### Application of ConDecon to Stereo-seq spatial transcriptomics data

We applied ConDecon to published Stereo-seq and single-cell RNA-seq data of the zebrafish embryo from Liu *et al*.^42^. The normalized gene expression data and cell type annotations were downloaded via h5ad files provided by the paper for the single-cell RNA-seq and Stereo-seq datasets of 3.3 hpf embryos. We filtered out genes expressed in < 3 cells, log-normalized gene counts, and identified variable genes using Scanpy^72^ (v1.9.1) (scanpy.pp.highly_variable_genes() function with default parameters). We used the top 3,350 variable genes to perform PCA and identified clusters with the Leiden algorithm at a resolution of 0.5 using the top 20 principal components. We computed diffusion pseudotime^43^ for the single-cell RNA-seq data using the scanpy.tl.dpt() function with default parameters, setting the cluster with highest expression of early blastodisc markers (otx1, dvl2, ctnnb1, axin1)^42^ as the root for the pseudotime calculation. We applied ConDecon with default parameters to infer single-cell abundances for each Stereo-seq pixel. We estimated the average pseudotime of each pixel using the approach described in subheading “Analysis of B-cell maturation”. To visualize the spatial cell differentiation trajectories derived from the inferred pseudotime spatial patterns, we computed the gradient over the pseudotime of each pixel using the immediately adjacent pixels. The resulting gradient vectors were smoothed using a gaussian kernel with standard deviation of 30 μm, truncated at 40 μm. We visualized the smoothed vector field using the matplotlib.pyplot.streamplot() function of Matplotlib (v3.5.3) with density = 1.5.

#### Application of ConDecon to ATAC-seq data

We applied ConDecon to published bulk and single-nucleus ATAC-seq data from two patient-derived melanoma cell lines (MM057 and MM087) profiled 0, 24, 48, or 72 hours after knocking down SOX10^50^. We used the 288 cells from MM087 as single-nucleus ATAC-seq reference data. To create a set of common peaks between the bulk and single-nucleus ATAC-seq data, we binned the genome into non-overlapping 10 kilobase bins. For each bin, we then aggregated the peaks that overlapped the bin. We assigned peaks that overlapped more than one bin to the bin with the smallest genome coordinates among the two overlapping bins. This resulted in 24,234 and 53,833 accessible bins in the single-nucleus and bulk data, respectively. To build a single-nucleus ATAC-seq data latent space, we used latent semantic indexing (LSI) as implemented in Signac^73^ (v1.1.0). We used bins that were open in at least 90% of the cells for term frequency-inverse document frequency (TF-IDF) normalization followed by singular value decomposition (SVD). We neglected the first component and visualized components 2 to 20 in two dimensions using UMAP. We applied ConDecon (with parameters max.center = 1, sigma_min_cells = 30, and sigma_max_cells = 75) to the bulk and single-nucleus bin-count matrices, using dimensions 2 to 11 of the latent space and the top 90% most variable bins (n = 22,362). We then estimated the average sampling time of the cells in each bulk sample based on the single-cell probabilities 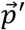 inferred by ConDecon (see subheading “Analysis of B-cell maturation”) and tested the association between the estimated average sampling time of the cells and the actual sampling time using Pearson’s correlation.

#### Gene expression deconvolution of ependymoma RNA-seq data

We downloaded the processed bulk (42 tumors) and single-nucleus RNA-seq data of pediatric ependymal tumors from Aubin *et al*.^21^ and applied ConDecon with default parameters using the top 5 latent dimensions and 5,000 variable genes. We assigned a score to each tumor cell in the single-nucleus data to represent its stage in the neuroepithelial-like to mesenchymal-like cell state transition. For that purpose, we aggregated for each cell the normalized expression values of the genes that are differentially expressed (FDR < 0.05, log fold-change > 3) between the clusters of neuroepithelial- and mesenchymal-like tumor cells. Similarly, we assigned a score representing the transition from a basal into a DAM state to each microglia in the single-nucleus data by aggregating the normalized expression values of the genes belonging to the DAM gene expression signature of Butovsky and Weiner^25^ (c.f. Fig. 2a in that reference). We then estimated the average scores of the tumor and microglial cells in each bulk sample using the same approach described above based on the inferred probabilities 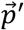 of ConDecon (subheading “Analysis of B cell maturation”). We tested the association between the estimated average microglial and mesenchymal scores of each bulk sample using Pearson’s correlation.

#### Gene expression deconvolution of spatial transcriptomic data of ependymal tumors

We downloaded the Visium spatial transcriptomic data of three pediatric posterior fossa ependymal tumors from Fu *et al*.^22^ (patients 459, 812, and 821). We used the regularized negative binomial regression model implemented in Seurat^74^ with default parameters to normalize the spatial transcriptomics gene counts. We applied ConDecon with default parameters to estimate the single-cell abundances of each spot in the spatial transcriptomic data, using the top 5 latent dimensions and 2,000 variable genes of the single-nucleus RNA-seq data of pediatric ependymal tumors from Aubin *et al*.^21^ as a reference. We then estimated the average DAM score of the tumor microglial cells in each spot based on the inferred probabilities 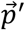 of ConDecon (subheading “Gene expression deconvolution of ependymoma RNA-seq data”). To visualize the spatial microglia differentiation trajectories, we computed the gradient over the DAM score of each pixel using the immediately adjacent pixels. The resulting gradient vectors were smoothed using a gaussian kernel with standard deviation of 300 μm, truncated at 400 μm.

#### Immunohistochemistry of ependymal tumors

De-identified formalin-fixed paraffin-embedded (FFPE) 5 μm tissue sections from one primary (7316-509) and one metastatic (7316-490, cortical metastasis) ependymal tumors located in or derived from the posterior fossa were provided by the Children’s Brain Tumor Tissue Network (CBTN) biorepository (Approved Biospecimen Project #29). The anatomic location of the tumors and their diagnosis were obtained from the surgical, radiology, and pathology reports. All the tissue sections and data were provided by the CBTN in a deidentified form according to the U.S. Department of Health and Human Services regulations and were not considered as Human Subjects Research by the Institutional Review Board of the University of Pennsylvania. Tissue handling procedures were performed according to the institutional regulations of the University of Pennsylvania and the Children’s Hospital of Philadelphia (CHOP). Adjacent FFPE sections from each of the tumors were stained with anti-CA9 (Novus, NB100-417SS), anti-IBA1 (Wako, 019-19741), and anti-GPNMB (R&D Systems, AF2550). Staining was performed on a Bond Max automated staining system (Leica Biosystems). The Bond Refine polymer staining kit (Leica Biosystems DS9800) was used for anti-IBA1 and anti-CA9. The Intense-R staining kit (Leica Biosystems, DS9263) was used for anti-GPNMB. The standard protocols were followed except for the primary antibody incubation, which was extended to 1 hour at room temperature. Antibodies were used at the following dilutions: anti-IBA1 1:2,000, anti-CA9 1:1,000, anti-GPNMB 1:500. Antigen retrieval was performed with E2 (anti-IBA1) or E1 (anti-CA9, anti-GPNMB) (Leica Microsystems) retrieval solution for 20 min. Slides were rinsed, dehydrated through ascending concentrations of ethanol and xylene, then cover-slipped. Stained slides were digitally scanned at 20x magnification on an Aperio AT2 slide scanner (Leica Biosystems).

### Quantification and statistical analysis

Two-sided Pearson correlation test of association was used in Figures 2B, 2D, 3A, 3B, 3C, 4C, 5G, 6D, S2, S3, S4, and S5. Two-sided Wilcoxon rank-sum test was used in Figures 4E, 5D, S1A, and S3D. One-sided Wilcoxon rank-sum test was used in Figure S3C. P values and sample sizes for each statistical test are described in the respective figure legend.

## Box 1

**Using gene ranks to infer cell abundances**

Given the gene expression profile of a bulk tissue, our goal is to infer the point in the space of cell abundance distributions over a reference single-cell RNA-seq dataset (of the same tissue type but possibly involving different cell abundances) that most closely represents the query bulk tissue. To reduce the effect of technical differences between single-cell RNA-seq and bulk RNA-seq measurements, we utilize gene ranks to compare the gene expression profile that results from aggregating single-cell gene expression levels across cells with the gene expression profile of the bulk tissue (Figure S1A). While different cell abundance configurations can lead to the same vector of gene ranks, this concern can be safely disregarded when working with single-cell datasets consisting of hundreds to thousands of variable genes.

Consider a reference single-cell dataset consisting of *J* cells and *T*variable genes and let *G* be the *T*× *J* expression table. The space of possible relative cell abundances in a synthetic bulk tissue constructed by sampling cells from the reference single-cell dataset consists of a (*J* − 1)-simplex, since cell proportions must add to 1. For example, in the case of 3 cells, the space of cell abundances consists of a triangle with unit-length sides, as illustrated below. For each point in the space of cell abundances, we can form a synthetic bulk RNA-seq dataset by aggregating the columns of *G* using weights given by the relative cell abundance of each cell. A bulk RNA-seq dataset can then be represented as a point in a *J*-dimensional space of gene rank correlations, where each dimension represents the value of the gene rank correlation distance of the bulk dataset with a cell in the reference single-cell dataset. Multiple points in the space of cell abundances may lead to the same point in the space of rank correlations, leading to a tessellation of the space of cell abundances. However, since the map between the space cell abundances and the space of gene rank correlations preserves local neighborhoods, we can think of the points in the space of gene rank correlations as a non-uniform pixelation of the space of cell abundances, where the resolution of the pixelation is controlled by the number of variable genes. Thus, for a sufficiently large number of variable genes (see Methods), it is possible to infer the relative cell abundances associated with a bulk RNA-seq dataset with high accuracy based on the gene rank correlations with the cells in a reference single-cell dataset, as shown below for a toy example consisting of 3 cells. The algorithm ConDecon tries to learn the map between the space of cell abundances and the space of gene rank correlations to infer cell abundances from bulk datasets.

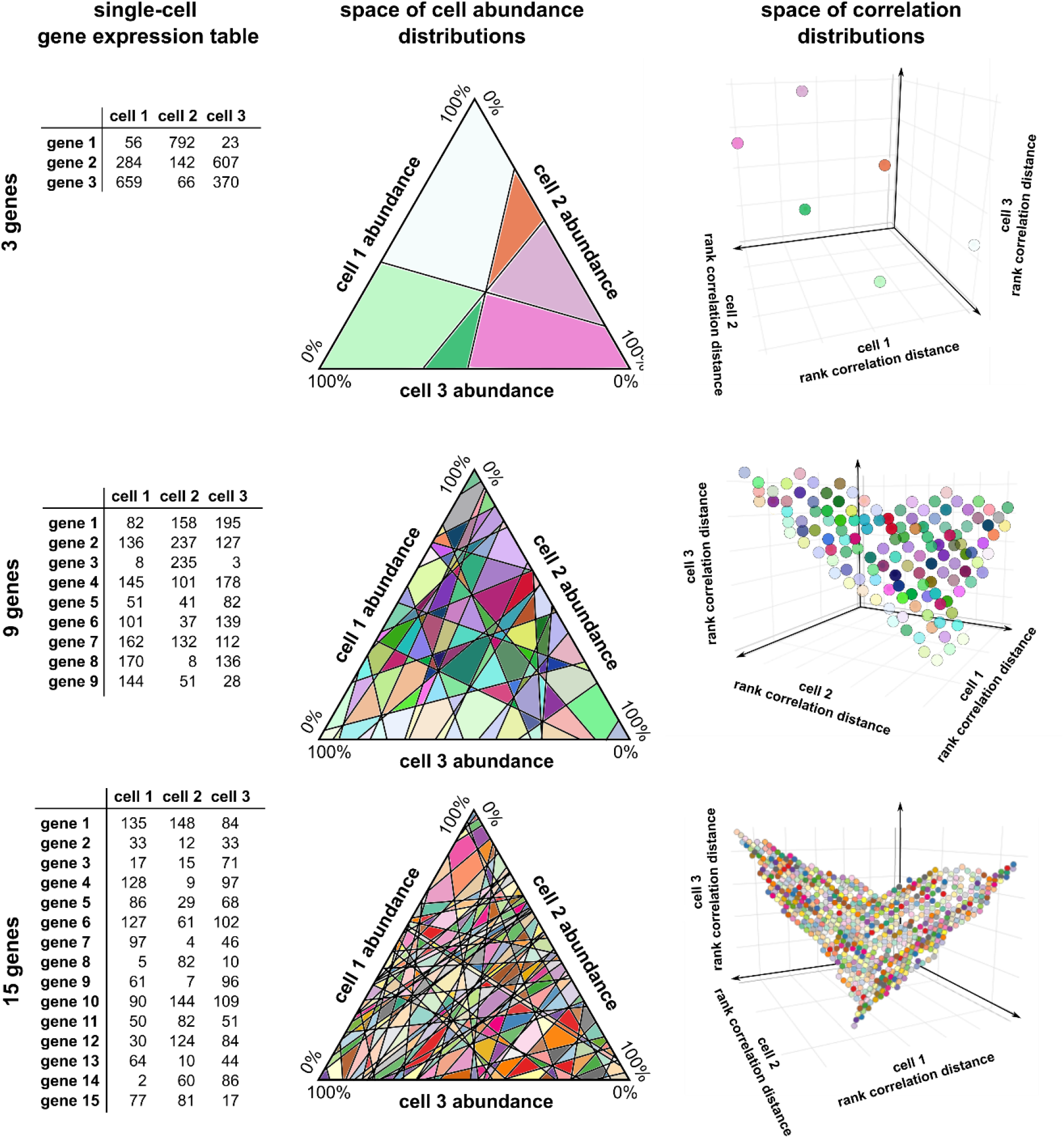

## References

1. Ramón, S. (1899). Textura del sistema nervioso del hombre y de los vertebrados: estudios sobre el plan estructural y composición histológica de los centros nerviosos adicionados de consideraciones fisiológicas fundadas en los nuevos descubrimientos (Moya).

2. Fawcett, D.W. (1969). An atlas of fine structure: The cell, its organelles and inclusions (Saunders).

3. Regev, A., Teichmann, S.A., Lander, E.S., Amit, I., Benoist, C., Birney, E., Bodenmiller, B., Campbell, P., Carninci, P., Clatworthy, M., et al. (2017). The Human Cell Atlas. Elife 6. 10.7554/eLife.27041.

4. Kolodziejczyk, A.A., Kim, J.K., Svensson, V., Marioni, J.C., and Teichmann, S.A. (2015). The technology and biology of single-cell RNA sequencing. Mol Cell 58, 610–620. 10.1016/j.molcel.2015.04.005.

5. Hwang, B., Lee, J.H., and Bang, D. (2018). Single-cell RNA sequencing technologies and bioinformatics pipelines. Exp Mol Med 50, 1–14. 10.1038/s12276-018-0071-8.

6. Denisenko, E., Guo, B.B., Jones, M., Hou, R., de Kock, L., Lassmann, T., Poppe, D., Clement, O., Simmons, R.K., Lister, R., and Forrest, A.R.R. (2020). Systematic assessment of tissue dissociation and storage biases in single-cell and single-nucleus RNA-seq workflows. Genome biology 21, 130. 10.1186/s13059-020-02048-6.

7. Avila Cobos, F., Vandesompele, J., Mestdagh, P., and De Preter, K. (2018). Computational deconvolution of transcriptomics data from mixed cell populations. Bioinformatics 34, 1969–1979. 10.1093/bioinformatics/bty019.

8. Mohammadi, S., Zuckerman, N., Goldsmith, A., and Grama, A. (2016). A critical survey of deconvolution methods for separating cell types in complex tissues. Proceedings of the IEEE 105, 340–366.

9. Newman, A.M., Liu, C.L., Green, M.R., Gentles, A.J., Feng, W., Xu, Y., Hoang, C.D., Diehn, M., and Alizadeh, A.A. (2015). Robust enumeration of cell subsets from tissue expression profiles. Nat Methods 12, 453–457. 10.1038/nmeth.3337.

10. Newman, A.M., Steen, C.B., Liu, C.L., Gentles, A.J., Chaudhuri, A.A., Scherer, F., Khodadoust, M.S., Esfahani, M.S., Luca, B.A., Steiner, D., et al. (2019). Determining cell type abundance and expression from bulk tissues with digital cytometry. Nat Biotechnol 37, 773–782. 10.1038/s41587-019-0114-2.

11. Baron, M., Veres, A., Wolock, S.L., Faust, A.L., Gaujoux, R., Vetere, A., Ryu, J.H., Wagner, B.K., Shen-Orr, S.S., Klein, A.M., et al. (2016). A Single-Cell Transcriptomic Map of the Human and Mouse Pancreas Reveals Inter- and Intra-cell Population Structure. Cell systems 3, 346–360 e344. 10.1016/j.cels.2016.08.011.

12. Wang, X., Park, J., Susztak, K., Zhang, N.R., and Li, M. (2019). Bulk tissue cell type deconvolution with multi-subject single-cell expression reference. Nature communications 10, 380. 10.1038/s41467-018-08023-x.

13. Hao, Y., Yan, M., Heath, B.R., Lei, Y.L., and Xie, Y. (2019). Fast and robust deconvolution of tumor infiltrating lymphocyte from expression profiles using least trimmed squares. PLoS computational biology 15, e1006976. 10.1371/journal.pcbi.1006976.

14. Racle, J., de Jonge, K., Baumgaertner, P., Speiser, D.E., and Gfeller, D. (2017). Simultaneous enumeration of cancer and immune cell types from bulk tumor gene expression data. Elife 6. 10.7554/eLife.26476.

15. Liebner, D.A., Huang, K., and Parvin, J.D. (2014). MMAD: microarray microdissection with analysis of differences is a computational tool for deconvoluting cell type-specific contributions from tissue samples. Bioinformatics 30, 682–689. 10.1093/bioinformatics/btt566.

16. Altboum, Z., Steuerman, Y., David, E., Barnett-Itzhaki, Z., Valadarsky, L., Keren-Shaul, H., Meningher, T., Mendelson, E., Mandelboim, M., Gat-Viks, I., and Amit, I. (2014). Digital cell quantification identifies global immune cell dynamics during influenza infection. Mol Syst Biol 10, 720. 10.1002/msb.134947.

17. Dong, M., Thennavan, A., Urrutia, E., Li, Y., Perou, C.M., Zou, F., and Jiang, Y. (2021). SCDC: bulk gene expression deconvolution by multiple single-cell RNA sequencing references. Briefings in bioinformatics 22, 416–427. 10.1093/bib/bbz166.

18. Jin, H., and Liu, Z. (2021). A benchmark for RNA-seq deconvolution analysis under dynamic testing environments. Genome biology 22, 102. 10.1186/s13059-021-02290-6.

19. Sturm, G., Finotello, F., Petitprez, F., Zhang, J.D., Baumbach, J., Fridman, W.H., List, M., and Aneichyk, T. (2019). Comprehensive evaluation of transcriptome-based cell-type quantification methods for immunooncology. Bioinformatics 35, i436–i445. 10.1093/bioinformatics/btz363.

20. Vallania, F., Tam, A., Lofgren, S., Schaffert, S., Azad, T.D., Bongen, E., Haynes, W., Alsup, M., Alonso, M., Davis, M., et al. (2018). Leveraging heterogeneity across multiple datasets increases cell-mixture deconvolution accuracy and reduces biological and technical biases. Nature communications 9, 4735. 10.1038/s41467-018-07242-6.

21. Aubin, R.G., Troisi, E.C., Montelongo, J., Alghalith, A.N., Nasrallah, M.P., Santi, M., and Camara, P.G. (2022). Pro-inflammatory cytokines mediate the epithelial-to-mesenchymal-like transition of pediatric posterior fossa ependymoma. Nature communications 13, 3936. 10.1038/s41467-022-31683-9.

22. Fu, R., Norris, G.A., Willard, N., Griesinger, A.M., Riemondy, K.A., Amani, V., Grimaldo, E., Harris, F., Hankinson, T.C., Mitra, S., et al. (2023). Spatial transcriptomic analysis delineates epithelial and mesenchymal subpopulations and transition stages in childhood ependymoma. Neuro Oncol 25, 786–798. 10.1093/neuonc/noac219.

23. Krasemann, S., Madore, C., Cialic, R., Baufeld, C., Calcagno, N., El Fatimy, R., Beckers, L., O’Loughlin, E., Xu, Y., Fanek, Z., et al. (2017). The TREM2-APOE Pathway Drives the Transcriptional Phenotype of Dysfunctional Microglia in Neurodegenerative Diseases. Immunity 47, 566–581 e569. 10.1016/j.immuni.2017.08.008.

24. Keren-Shaul, H., Spinrad, A., Weiner, A., Matcovitch-Natan, O., Dvir-Szternfeld, R., Ulland, T.K., David, E., Baruch, K., Lara-Astaiso, D., Toth, B., et al. (2017). A Unique Microglia Type Associated with Restricting Development of Alzheimer’s Disease. Cell 169, 1276–1290 e1217. 10.1016/j.cell.2017.05.018.

25. Butovsky, O., and Weiner, H.L. (2018). Microglial signatures and their role in health and disease. Nat Rev Neurosci 19, 622–635. 10.1038/s41583-018-0057-5.

26. Sun, D., Guan, X., Moran, A.E., Wu, L.Y., Qian, D.Z., Schedin, P., Dai, M.S., Danilov, A.V., Alumkal, J.J., Adey, A.C., et al. (2022). Identifying phenotype-associated subpopulations by integrating bulk and single-cell sequencing data. Nat Biotechnol 40, 527–538. 10.1038/s41587-021-01091-3.

27. Zappia, L., Phipson, B., and Oshlack, A. (2017). Splatter: simulation of single-cell RNA sequencing data. Genome biology 18, 174. 10.1186/s13059-017-1305-0.

28. Oetjen, K.A., Lindblad, K.E., Goswami, M., Gui, G., Dagur, P.K., Lai, C., Dillon, L.W., McCoy, J.P., and Hourigan, C.S. (2018). Human bone marrow assessment by single-cell RNA sequencing, mass cytometry, and flow cytometry. JCI Insight 3. 10.1172/jci.insight.124928.

29. Tabula Muris, C. (2020). A single-cell transcriptomic atlas characterizes ageing tissues in the mouse. Nature 583, 590–595. 10.1038/s41586-020-2496-1.

30. Regev, A., Teichmann, S., Lander, E.S., Amit, I., Benoist, C., Birney, E., Bodenmiller, B., Campbell, P., Carninci, P., and Clatworthy, M. (2017). The Human Cell Atlas. bioRxiv, 121202.

31. Rozenblatt-Rosen, O., Regev, A., Oberdoerffer, P., Nawy, T., Hupalowska, A., Rood, J.E., Ashenberg, O., Cerami, E., Coffey, R.J., Demir, E., et al. (2020). The Human Tumor Atlas Network: Charting Tumor Transitions across Space and Time at Single-Cell Resolution. Cell 181, 236–249. 10.1016/j.cell.2020.03.053.

32. Avila Cobos, F., Alquicira-Hernandez, J., Powell, J.E., Mestdagh, P., and De Preter, K. (2020). Benchmarking of cell type deconvolution pipelines for transcriptomics data. Nature communications 11, 5650. 10.1038/s41467-020-19015-1.

33. Enge, M., Arda, H.E., Mignardi, M., Beausang, J., Bottino, R., Kim, S.K., and Quake, S.R. (2017). Single-Cell Analysis of Human Pancreas Reveals Transcriptional Signatures of Aging and Somatic Mutation Patterns. Cell 171, 321–330 e314. 10.1016/j.cell.2017.09.004.

34. Han, X., Zhou, Z., Fei, L., Sun, H., Wang, R., Chen, Y., Chen, H., Wang, J., Tang, H., Ge, W., et al. (2020). Construction of a human cell landscape at single-cell level. Nature 581, 303–309. 10.1038/s41586-020-2157-4.

35. Jew, B., Alvarez, M., Rahmani, E., Miao, Z., Ko, A., Garske, K.M., Sul, J.H., Pietilainen, K.H., Pajukanta, P., and Halperin, E. (2020). Accurate estimation of cell composition in bulk expression through robust integration of single-cell information. Nature communications 11, 1971. 10.1038/s41467-020-15816-6.

36. Lab, A.s.L.S.F.C.C.D. (2023). The Single-cell Pediatric Cancer Data Lab, https://scpca.alexslemonade.org/.

37. Frishberg, A., Peshes-Yaloz, N., Cohn, O., Rosentul, D., Steuerman, Y., Valadarsky, L., Yankovitz, G., Mandelboim, M., Iraqi, F.A., Amit, I., et al. (2019). Cell composition analysis of bulk genomics using singlecell data. Nat Methods 16, 327–332. 10.1038/s41592-019-0355-5.

38. Johnson, K.M., Owen, K., and Witte, P.L. (2002). Aging and developmental transitions in the B cell lineage. Int Immunol 14, 1313–1323. 10.1093/intimm/dxf092.

39. Chen, L., Wang, J., Liu, J., Wang, H., Hillyer, C.D., Blanc, L., An, X., and Mohandas, N. (2021). Dynamic changes in murine erythropoiesis from birth to adulthood: implications for the study of murine models of anemia. Blood Adv 5, 16–25. 10.1182/bloodadvances.2020003632.

40. Chiossone, L., Chaix, J., Fuseri, N., Roth, C., Vivier, E., and Walzer, T. (2009). Maturation of mouse NK cells is a 4-stage developmental program. Blood 113, 5488–5496. 10.1182/blood-2008-10-187179.

41. Song, L., Sun, X., Qi, T., and Yang, J. (2023). Mixed model-based deconvolution of cell-state abundances (MeDuSA) along a one-dimensional trajectory. Nat Comput Sci 3, 630–643. 10.1038/s43588-023-00487-2.

42. Liu, C., Li, R., Li, Y., Lin, X., Zhao, K., Liu, Q., Wang, S., Yang, X., Shi, X., Ma, Y., et al. (2022). Spatiotemporal mapping of gene expression landscapes and developmental trajectories during zebrafish embryogenesis. Dev Cell 57, 1284–1298 e1285. 10.1016/j.devcel.2022.04.009.

43. Haghverdi, L., Buttner, M., Wolf, F.A., Buettner, F., and Theis, F.J. (2016). Diffusion pseudotime robustly reconstructs lineage branching. Nat Methods 13, 845–848. 10.1038/nmeth.3971.

44. Kimmel, C.B., Ballard, W.W., Kimmel, S.R., Ullmann, B., and Schilling, T.F. (1995). Stages of embryonic development of the zebrafish. Dev Dyn 203, 253–310. 10.1002/aja.1002030302.

45. Danaher, P., Kim, Y., Nelson, B., Griswold, M., Yang, Z., Piazza, E., and Beechem, J.M. (2022). Advances in mixed cell deconvolution enable quantification of cell types in spatial transcriptomic data. Nature communications 13, 385. 10.1038/s41467-022-28020-5.

46. Dong, R., and Yuan, G.C. (2021). SpatialDWLS: accurate deconvolution of spatial transcriptomic data. Genome biology 22, 145. 10.1186/s13059-021-02362-7.

47. Elosua-Bayes, M., Nieto, P., Mereu, E., Gut, I., and Heyn, H. (2021). SPOTlight: seeded NMF regression to deconvolute spatial transcriptomics spots with single-cell transcriptomes. Nucleic acids research 49, e50. 10.1093/nar/gkab043.

48. Ma, Y., and Zhou, X. (2022). Spatially informed cell-type deconvolution for spatial transcriptomics. Nat Biotechnol 40, 1349–1359. 10.1038/s41587-022-01273-7.

49. Song, Q., and Su, J. (2021). DSTG: deconvoluting spatial transcriptomics data through graph-based artificial intelligence. Briefings in bioinformatics 22. 10.1093/bib/bbaa414.

50. Bravo Gonzalez-Blas, C., Minnoye, L., Papasokrati, D., Aibar, S., Hulselmans, G., Christiaens, V., Davie, K., Wouters, J., and Aerts, S. (2019). cisTopic: cis-regulatory topic modeling on single-cell ATAC-seq data. Nat Methods 16, 397–400. 10.1038/s41592-019-0367-1.

51. Junger, S.T., Timmermann, B., and Pietsch, T. (2021). Pediatric ependymoma: an overview of a complex disease. Childs Nerv Syst 37, 2451–2463. 10.1007/s00381-021-05207-7.

52. Saleh, A.H., Samuel, N., Juraschka, K., Saleh, M.H., Taylor, M.D., and Fehlings, M.G. (2022). The biology of ependymomas and emerging novel therapies. Nat Rev Cancer 22, 208–222. 10.1038/s41568-021-00433-2.

53. Wu, J., Armstrong, T.S., and Gilbert, M.R. (2016). Biology and management of ependymomas. Neuro Oncol 18, 902–913. 10.1093/neuonc/now016.

54. Gillen, A.E., Riemondy, K.A., Amani, V., Griesinger, A.M., Gilani, A., Venkataraman, S., Madhavan, K., Prince, E., Sanford, B., Hankinson, T.C., et al. (2020). Single-Cell RNA Sequencing of Childhood Ependymoma Reveals Neoplastic Cell Subpopulations That Impact Molecular Classification and Etiology. Cell Rep 32, 108023. 10.1016/j.celrep.2020.108023.

55. Gojo, J., Englinger, B., Jiang, L., Hubner, J.M., Shaw, M.L., Hack, O.A., Madlener, S., Kirchhofer, D., Liu, I., Pyrdol, J., et al. (2020). Single-Cell RNA-Seq Reveals Cellular Hierarchies and Impaired Developmental Trajectories in Pediatric Ependymoma. Cancer Cell 38, 44–59 e49. 10.1016/j.ccell.2020.06.004.

56. Bergen, V., Lange, M., Peidli, S., Wolf, F.A., and Theis, F.J. (2020). Generalizing RNA velocity to transient cell states through dynamical modeling. Nat Biotechnol. 10.1038/s41587-020-0591-3.

57. La Manno, G., Soldatov, R., Zeisel, A., Braun, E., Hochgerner, H., Petukhov, V., Lidschreiber, K., Kastriti, M.E., Lonnerberg, P., Furlan, A., et al. (2018). RNA velocity of single cells. Nature 560, 494–498. 10.1038/s41586-018-0414-6.

58. Preusser, M., Wolfsberger, S., Haberler, C., Breitschopf, H., Czech, T., Slavc, I., Harris, A.L., Acker, T., Budka, H., and Hainfellner, J.A. (2005). Vascularization and expression of hypoxia-related tissue factors in intracranial ependymoma and their impact on patient survival. Acta Neuropathol 109, 211–216. 10.1007/s00401-004-0938-8.

59. Fu, R., Norris, G.A., Willard, N., Griesinger, A.M., Riemondy, K.A., Amani, V., Grimaldo, E., Harris, F., Hankinson, T.C., Mitra, S., et al. (2022). Spatial transcriptomic analysis delineates epithelial and mesenchymal subpopulations and transition stages in childhood ependymoma. Neuro Oncol. 10.1093/neuonc/noac219.

60. Xiong, A., Zhang, J., Chen, Y., Zhang, Y., and Yang, F. (2022). Integrated single-cell transcriptomic analyses reveal that GPNMB-high macrophages promote PN-MES transition and impede T cell activation in GBM. EBioMedicine 83, 104239. 10.1016/j.ebiom.2022.104239.

61. Satija, R., Farrell, J.A., Gennert, D., Schier, A.F., and Regev, A. (2015). Spatial reconstruction of single-cell gene expression data. Nat Biotechnol 33, 495–502. 10.1038/nbt.3192.

62. Dietrich, A., Sturm, G., Merotto, L., Marini, F., Finotello, F., and List, M. (2022). SimBu: bias-aware simulation of bulk RNA-seq data with variable cell-type composition. Bioinformatics 38, ii141–ii147. 10.1093/bioinformatics/btac499.

63. Hippen, A.A., Omran, D.K., Weber, L.M., Jung, E., Drapkin, R., Doherty, J.A., Hicks, S.C., and Greene, C.S. (2023). Performance of computational algorithms to deconvolve heterogeneous bulk ovarian tumor tissue depends on experimental factors. Genome biology 24, 239. 10.1186/s13059-023-03077-7.

64. Kendall, M.G. (1938). A new measure of rank correlation. Biometrika 30, 81–93.

65. Diaconis, P. (1988). Group representations in probability and statistics. Lecture notes-monograph series 11, i–192.

66. Petersen, P. (2006). Riemannian geometry (Springer).

67. Kendall, M.G., and Gibbons, J.D. (1990). Rank Correlation Methods (Edward Arnold).

68. Lun, A.T., McCarthy, D.J., and Marioni, J.C. (2016). A step-by-step workflow for low-level analysis of singlecell RNA-seq data with Bioconductor. F1000Res 5, 2122. 10.12688/f1000research.9501.2.

69. Segerstolpe, A., Palasantza, A., Eliasson, P., Andersson, E.M., Andreasson, A.C., Sun, X., Picelli, S., Sabirsh, A., Clausen, M., Bjursell, M.K., et al. (2016). Single-Cell Transcriptome Profiling of Human Pancreatic Islets in Health and Type 2 Diabetes. Cell Metab 24, 593–607. 10.1016/j.cmet.2016.08.020.

70. Nestorowa, S., Hamey, F.K., Pijuan Sala, B., Diamanti, E., Shepherd, M., Laurenti, E., Wilson, N.K., Kent, D.G., and Gottgens, B. (2016). A single-cell resolution map of mouse hematopoietic stem and progenitor cell differentiation. Blood 128, e20–31. 10.1182/blood-2016-05-716480.

71. Cao, J., Spielmann, M., Qiu, X., Huang, X., Ibrahim, D.M., Hill, A.J., Zhang, F., Mundlos, S., Christiansen, L., Steemers, F.J., et al. (2019). The single-cell transcriptional landscape of mammalian organogenesis. Nature 566, 496–502. 10.1038/s41586-019-0969-x.

72. Wolf, F.A., Angerer, P., and Theis, F.J. (2018). SCANPY: large-scale single-cell gene expression data analysis. Genome biology 19, 15. 10.1186/s13059-017-1382-0.

73. Stuart, T., Srivastava, A., Madad, S., Lareau, C.A., and Satija, R. (2021). Single-cell chromatin state analysis with Signac. Nat Methods 18, 1333–1341. 10.1038/s41592-021-01282-5.

74. Hafemeister, C., and Satija, R. (2019). Normalization and variance stabilization of single-cell RNA-seq data using regularized negative binomial regression. Genome biology 20, 296. 10.1186/s13059-019-1874-1.

