## Supplemental Figures for "Clustering-independent estimation of cell abundances in bulk tissues using single-cell RNA-seq data"

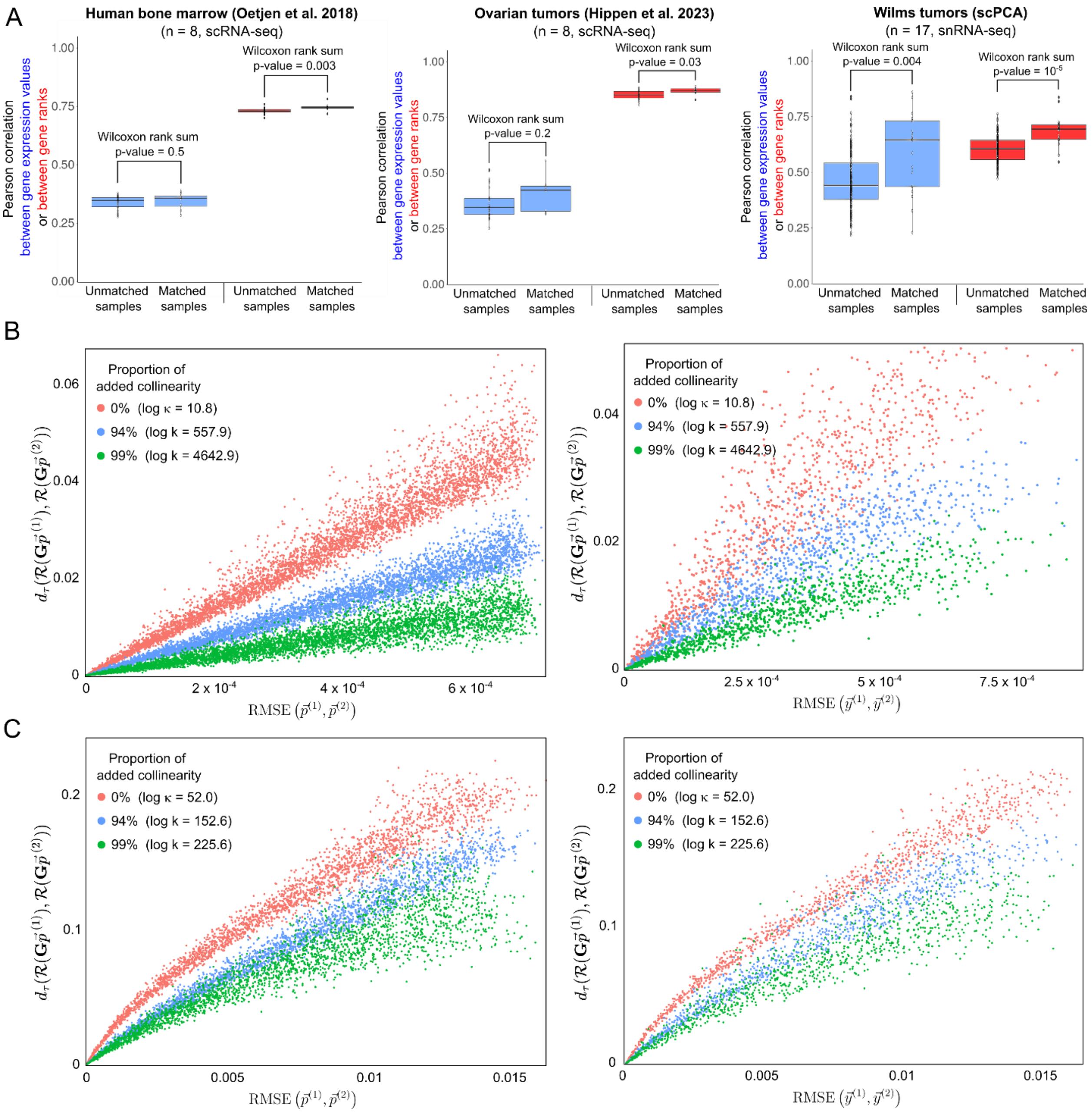


**Figure S1. Using gene ranks to infer cell abundance from gene expression data of bulk tissues. (A)** Pearson correlation coefficient between the expression levels or between the gene ranks for 3 datasets consisting of single-cell/nucleus and bulk RNA-seq data from the same samples, encompassing human bone marrow[1], high-grade serous ovarian tumors[2], and Wilms tumors from the single-cell pediatric cancer atlas (scPCA). For each sample, we aggregated the single cell or single nucleus counts across the cells to construct a synthetic bulk RNA-seq dataset for the sample. We then computed the Pearson correlation coefficient between the expression levels, or between the gene ranks, of the synthetic and actual bulk RNA-seq datasets using the 2,000 most variable genes. This analysis shows gene ranks from aggregated single-cell/nucleus can better discriminate between samples than aggregated gene expression values. In particular, the concordance between the ranks of the genes is higher than between the gene expression values, and the correlation between gene ranks in synthetic and real bulk datasets from the same individual ('matched samples') is significantly higher than in synthetic and real bulk datasets from different individuals ('unmatched samples'). **(B)** Distance (Kendall's $\tau$ distance, $d_\tau$) in the space of gene rank correlation distributions as a function of the distance in the space of cell abundance distributions (root mean square error, RMSE) for various levels of collinearity in the gene expression matrix $\boldsymbol{G}$. Each point corresponds to a pair of simulated random cell abundance distributions over 8,000 cells from a human bone-marrow single-cell RNA-seq dataset[1]. The bulk gene expression profile corresponding to each simulated distribution is obtained by aggregating the single-cell gene expression counts of the individual cells (left) or top 10 principal components (right) according to their probability for the 2,000 most variable genes. Additional collinearity is included by replacing the gene expression profile of a fraction of the cells with rescaled copies of the expression profile of other cells in the single-cell dataset. The amount of added collinearity and the logarithm of the resulting condition number for the gene expression matrix ($\log \kappa$) are indicated. As predicted from the mathematical foundation of ConDecon, for a sufficiently large number of variable genes, the distance between two bulk datasets in the space of rank correlations is small ($d_\tau\left(\mathcal{R}\left(\boldsymbol{G}\vec{p}^{(1)}\right), \mathcal{R}\left(\boldsymbol{G}\vec{p}^{(2)}\right)\right) \simeq 0$) if and only if their cell abundance composition is very similar ($\mathrm{RMSE}\left(\vec{p}^{(1)}, \vec{p}^{(2)}\right) \simeq 0$). **(C)** Same as in (B), but the simulated random cell abundance distributions are built by sampling from the probability simplex using beta distributions $\beta(a_1, a_2)$ and $\beta(a_2, a_1)$, with $a_1 = 0.5$ and $a_2 = 8$, instead of uniform sampling.

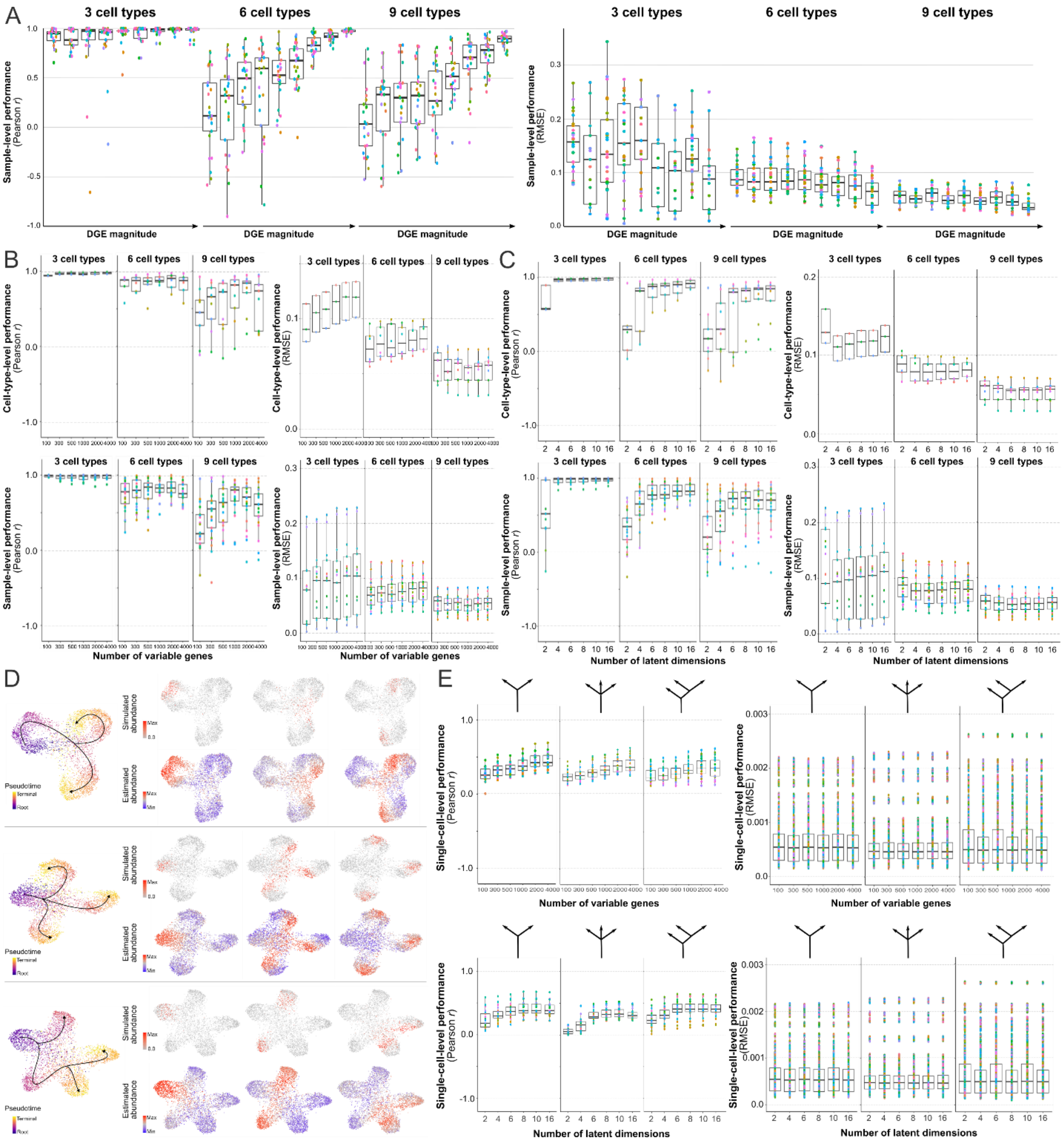


**Figure S2. Deconvolution of simulated bulk RNA-seq data of discrete cell populations and continuous cell differentiation processes. (A)** Sample-level Pearson correlation coefficient and root mean square error (RMSE) between simulated and estimated cell population abundances in simulations of bulk RNA-seq datasets (*n* = 675) with 3, 6, or 9 discrete cell populations and varying degree of differential gene expression (DGE). **(B, C)** Cell-type- and sample-level Pearson correlation coefficient and RMSE between simulated and estimated cell population abundances in simulated bulk RNA-seq datasets ($n$ = 675) with 3, 6, or 9 discrete cell populations as a function of the number of variable genes (B) and latent dimensions (C) used by ConDecon. **(D)** Cell abundance estimation in 9 simulated bulk RNA-seq datasets of 3 cell differentiation processes with 1 precursor and 2 or 3 terminally differentiated cell states. Left: The UMAP representation of each simulated single-cell RNA-seq dataset is colored by the simulated pseudotime. Right: For each single-cell dataset, the simulated (top) and estimated (bottom) cell abundances are shown for 3 bulk RNA-seq datasets constructed by sampling cells non-uniformly from the single-cell dataset. **(E)** Single-cell-level Pearson correlation coefficient and RMSE between simulated and estimated cell abundances in simulations of bulk RNA-seq datasets with 3 different topologies as a function of the number of variable genes (top) and latent dimensions (bottom) used by ConDecon. The topologies of the cell differentiation processes are indicated at the top.

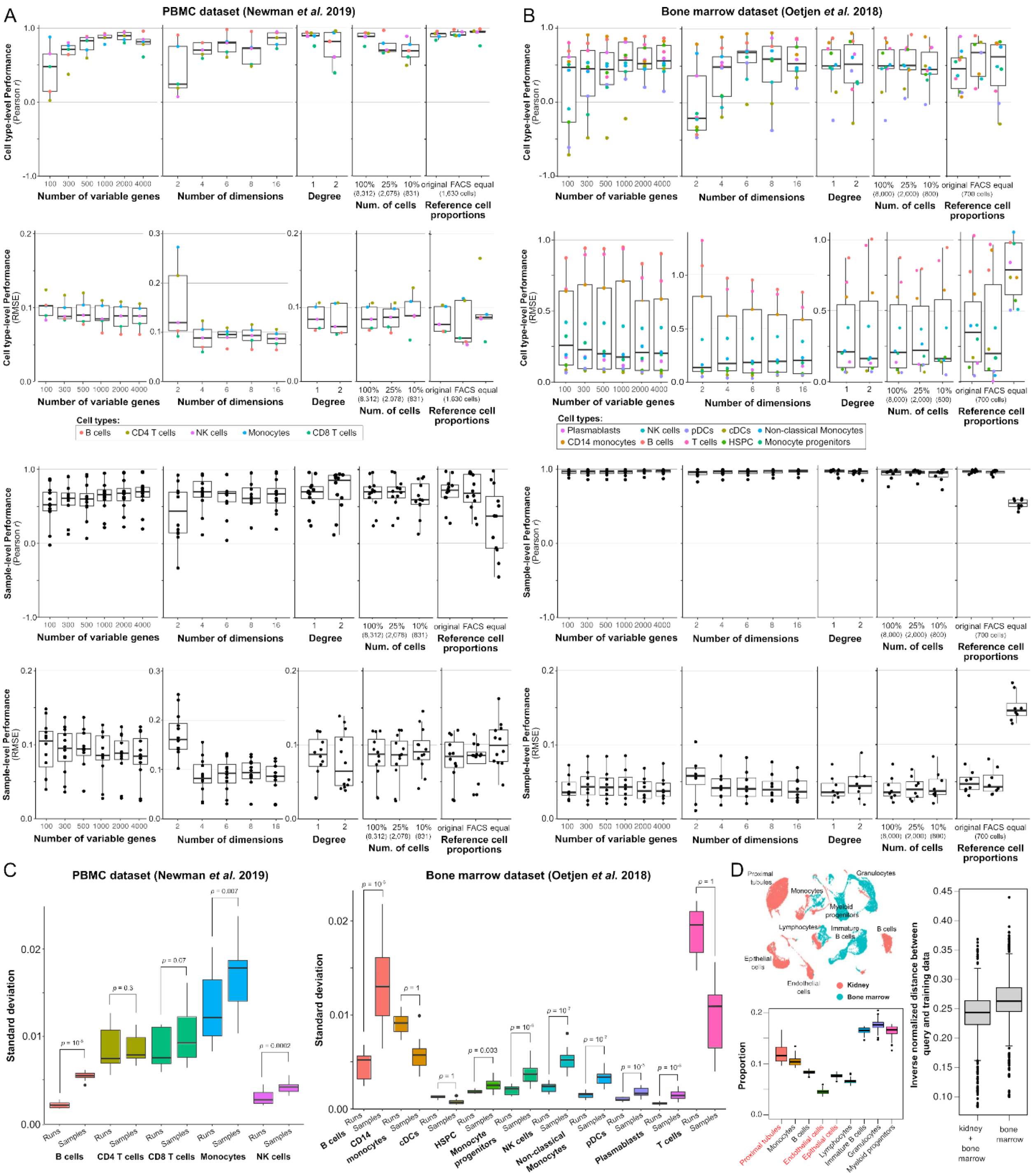

**Figure S3. Deconvolution of bulk RNA-seq data of PBMCs and bone marrow as a function of ConDecon's parameters. (A, B)** Single-cell-level Pearson correlation coefficient and root mean square error (RMSE) between the estimated cell abundances by ConDecon and the observed abundances by FACS across samples (cell-type-level performance) and cell types (sample-level performance) for bulk RNA-seq datasets consisting of 8 bone marrow[1] (A) and 12 PBMC[3] (B) samples as a function of the number of variable genes, the number of latent dimensions, the degree of the polynomial, the number of cells in the reference single-cell RNA-seq dataset, and the similitude between the cell proportions in the reference single-cell data set and the query bulk data, where “original”, “FACS”, and “equal” indicate the proportions in the original single-cell dataset, in the FACS data, or equal proportions for all the cell populations present in the reference single-cell dataset, respectively. **(C)** Standard deviation of the cell type abundance inferences of ConDecon across 20 different random initializations (“runs”) and across samples (“samples”). The variability of the inferred abundances across runs is significantly smaller than the variability across samples for almost all cell types. 1-sided Wilcoxon rank sum test *p*-values are indicated. **(D)** Deconvolution of mouse bone marrow bulk data using reference single-cell RNA-seq data of mouse bone marrow and kidney from the Tabula Muris Senis[4]. Left, top: UMAP representation of the combined kidney and bone marrow single-cell RNA-seq datasets. Left, bottom: Inferred cell type abundances for bulk RNA-seq data from 53 bone marrow samples. The inferences of ConDecon are affected by the large mismatch between the reference and query datasets, with 24% of the probability mass assigned to kidney-specific cell populations (indicated in red). Right: The inverse distance between the point that corresponds to the query bulk sample and the 10 nearest training data points in the space of probability distributions, normalized by the average distance between training data points, can be used as an indicator of the quality of the inferences made by ConDecon. The inverse distance varies between 0 (for single-cell reference data unrelated to the query bulk data) and approximately 1 (for single-cell reference data that accurately match the query bulk data). In the figure, the inverse distance is significantly increased when using a bone marrow instead of a combined bone marrow and kidney single-cell RNA-seq dataset to deconvolve the bone marrow bulk RNA-seq data (2-sided Wilcoxon rank-sum test *p*-value $< 10^{-16}$).

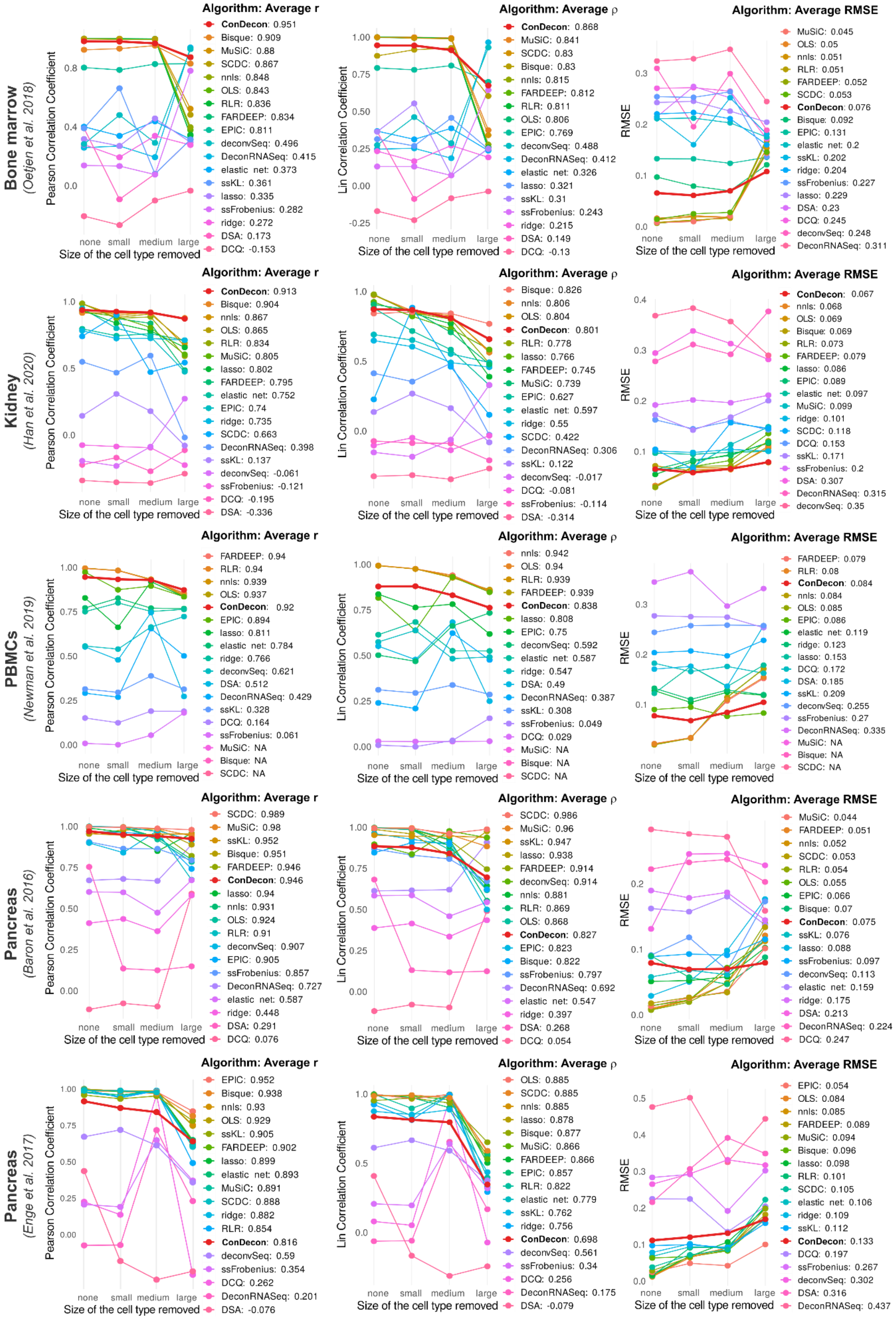
Bone marrow
(Oetjen et al. 2018)
Pearson Correlation Coefficient
Size of the cell type removed
none small medium large
Algorithm: Average r
ConDecon: 0.951
Bisque: 0.909
MuSiC: 0.88
SCDC: 0.867
nnls: 0.848
OLS: 0.843
RLR: 0.836
FARDEEP: 0.834
EPIC: 0.811
deconvSeq: 0.496
DeconRNASeq: 0.415
elastic net: 0.373
ssKL: 0.361
lasso: 0.335
ssFrobenius: 0.282
ridge: 0.272
DSA: 0.173
DCQ: -0.153
Lin Correlation Coefficient
Algorithm: Average ρ
ConDecon: 0.868
MuSiC: 0.841
SCDC: 0.83
Bisque: 0.83
nnls: 0.815
FARDEEP: 0.812
RLR: 0.811
OLS: 0.806
EPIC: 0.769
deconvSeq: 0.488
DeconRNASeq: 0.412
elastic net: 0.326
lasso: 0.321
ssKL: 0.31
ssFrobenius: 0.243
ridge: 0.215
DSA: 0.149
DCQ: -0.13
RMSE
Algorithm: Average RMSE
MuSiC: 0.045
OLS: 0.05
nnls: 0.051
RLR: 0.051
FARDEEP: 0.052
SCDC: 0.053
ConDecon: 0.076
Bisque: 0.092
EPIC: 0.131
elastic net: 0.2
ssKL: 0.202
ridge: 0.204
ssFrobenius: 0.227
lasso: 0.229
DSA: 0.23
DCQ: 0.245
deconvSeq: 0.248
DeconRNASeq: 0.311
Kidney
(Han et al. 2020)
Algorithm: Average r
ConDecon: 0.913
Bisque: 0.904
nnls: 0.867
OLS: 0.865
RLR: 0.834
MuSiC: 0.805
lasso: 0.802
FARDEEP: 0.795
elastic net: 0.752
EPIC: 0.74
ridge: 0.735
SCDC: 0.663
DeconRNASeq: 0.398
ssKL: 0.137
deconvSeq: -0.061
ssFrobenius: -0.121
DCQ: -0.195
DSA: -0.336
Algorithm: Average ρ
Bisque: 0.826
nnls: 0.806
OLS: 0.804
ConDecon: 0.801
RLR: 0.778
lasso: 0.766
FARDEEP: 0.745
MuSiC: 0.739
EPIC: 0.627
elastic net: 0.597
ridge: 0.55
SCDC: 0.422
DeconRNASeq: 0.306
ssKL: 0.122
deconvSeq: -0.017
DCQ: -0.081
ssFrobenius: -0.114
DSA: -0.314
Algorithm: Average RMSE
ConDecon: 0.067
nnls: 0.068
OLS: 0.069
Bisque: 0.069
RLR: 0.073
FARDEEP: 0.079
lasso: 0.086
EPIC: 0.089
elastic net: 0.097
MuSiC: 0.099
ridge: 0.101
SCDC: 0.118
DCQ: 0.153
ssKL: 0.171
ssFrobenius: 0.2
DSA: 0.307
DeconRNASeq: 0.315
deconvSeq: 0.35
PBMCs
(Newman et al. 2019)
Algorithm: Average r
FARDEEP: 0.94
RLR: 0.94
nnls: 0.939
OLS: 0.937
ConDecon: 0.92
EPIC: 0.894
lasso: 0.811
elastic net: 0.784
ridge: 0.766
deconvSeq: 0.621
DSA: 0.512
DeconRNASeq: 0.429
ssKL: 0.328
DCQ: 0.164
ssFrobenius: 0.061
MuSiC: NA
Bisque: NA
SCDC: NA
Algorithm: Average ρ
nnls: 0.942
OLS: 0.94
RLR: 0.939
FARDEEP: 0.939
ConDecon: 0.838
lasso: 0.808
EPIC: 0.75
deconvSeq: 0.592
elastic net: 0.587
ridge: 0.547
DSA: 0.49
DeconRNASeq: 0.387
ssKL: 0.308
ssFrobenius: 0.049
DCQ: 0.029
MuSiC: NA
Bisque: NA
SCDC: NA
Algorithm: Average RMSE
FARDEEP: 0.079
RLR: 0.08
ConDecon: 0.084
nnls: 0.084
OLS: 0.085
EPIC: 0.086
elastic net: 0.119
ridge: 0.123
lasso: 0.153
DCQ: 0.172
DSA: 0.185
ssKL: 0.209
deconvSeq: 0.255
ssFrobenius: 0.27
DeconRNASeq: 0.335
MuSiC: NA
Bisque: NA
SCDC: NA
Pancreas
(Baron et al. 2016)
Algorithm: Average r
SCDC: 0.989
MuSiC: 0.98
ssKL: 0.952
Bisque: 0.951
FARDEEP: 0.946
ConDecon: 0.946
lasso: 0.94
nnls: 0.931
OLS: 0.924
RLR: 0.91
deconvSeq: 0.907
EPIC: 0.905
ssFrobenius: 0.857
DeconRNASeq: 0.727
elastic net: 0.587
ridge: 0.448
DSA: 0.291
DCQ: 0.076
Algorithm: Average ρ
SCDC: 0.986
MuSiC: 0.96
ssKL: 0.947
lasso: 0.938
FARDEEP: 0.914
deconvSeq: 0.914
nnls: 0.881
RLR: 0.869
OLS: 0.868
ConDecon: 0.827
EPIC: 0.823
Bisque: 0.822
ssFrobenius: 0.797
DeconRNASeq: 0.692
elastic net: 0.547
ridge: 0.397
DSA: 0.268
DCQ: 0.054
Algorithm: Average RMSE
MuSiC: 0.044
FARDEEP: 0.051
nnls: 0.052
SCDC: 0.053
RLR: 0.054
OLS: 0.055
EPIC: 0.066
Bisque: 0.07
ConDecon: 0.075
ssKL: 0.076
lasso: 0.088
ssFrobenius: 0.097
deconvSeq: 0.113
elastic net: 0.159
ridge: 0.175
DSA: 0.213
DeconRNASeq: 0.224
DCQ: 0.247
Pancreas
(Enge et al. 2017)
Algorithm: Average r
EPIC: 0.952
Bisque: 0.938
nnls: 0.93
OLS: 0.929
ssKL: 0.905
FARDEEP: 0.902
lasso: 0.899
elastic net: 0.893
MuSiC: 0.891
SCDC: 0.888
ridge: 0.882
RLR: 0.854
ConDecon: 0.816
deconvSeq: 0.59
ssFrobenius: 0.354
DCQ: 0.262
DeconRNASeq: 0.201
DSA: -0.076
Algorithm: Average ρ
OLS: 0.885
SCDC: 0.885
nnls: 0.885
lasso: 0.878
Bisque: 0.877
MuSiC: 0.866
FARDEEP: 0.866
EPIC: 0.857
RLR: 0.822
elastic net: 0.779
ssKL: 0.762
ridge: 0.756
ConDecon: 0.698
deconvSeq: 0.561
ssFrobenius: 0.34
DCQ: 0.256
DeconRNASeq: 0.175
DSA: -0.079
Algorithm: Average RMSE
EPIC: 0.054
OLS: 0.084
nnls: 0.085
FARDEEP: 0.089
MuSiC: 0.094
Bisque: 0.096
lasso: 0.098
RLR: 0.101
SCDC: 0.105
elastic net: 0.106
ridge: 0.109
ssKL: 0.112
ConDecon: 0.133
DCQ: 0.197
ssFrobenius: 0.267
deconvSeq: 0.302
DSA: 0.316
DeconRNASeq: 0.437

**Figure S4. Benchmarking of the aggregated cell abundance estimates of ConDecon in comparison to seventeen other methods for gene expression deconvolution.** Evaluation of the aggregated cell abundance estimates of ConDecon and the cell type abundance estimates of 17 other deconvolution methods across 5 datasets using the benchmarking pipeline of Avila-Cobos *et al.* For each algorithm and dataset, the Pearson's correlation coefficient (left), the Lin's concordance correlation coefficient, and the root mean squared error (RMSE) (right) of the estimates, combined across samples and cell types, is shown for cases where there is none, one small, one medium, or one large cell population missing in the reference single-cell data.

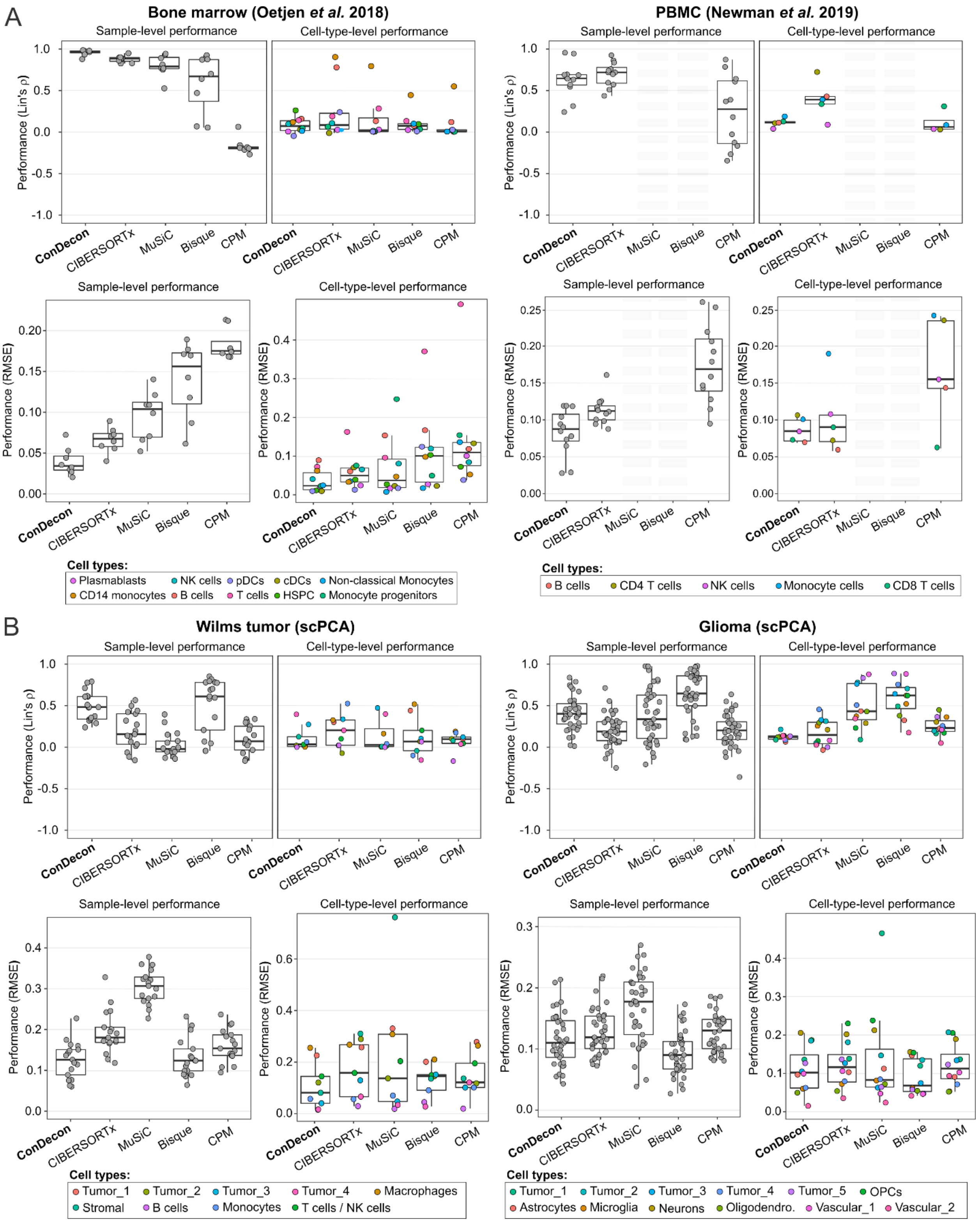


**Figure S5. Comparison between cell type abundance estimates derived from FACS and single-nucleus RNA-seq data and those from ConDecon and 4 other deconvolution**

**methods. (A, B)** Two bulk RNA-seq datasets consisting of 8 bone marrow[1] (A, left) and 12 PBMC[3] (A, right) samples, for which paired FACS data are available, as well as two bulk RNA-seq datasets consisting of 17 Wilms tumor (B, right) and 37 pediatric glioma (B, left) samples, for which paired single-nuclei RNA-seq data are available, were considered. The sample-level and cell-type-level RMSE and Lin's concordance correlation coefficient are shown for each algorithm in each dataset. We were unable to apply MuSiC and Bisque to the PBMC dataset since these methods require that the reference single-cell RNA-seq data consists of at least 2 biological replicates.

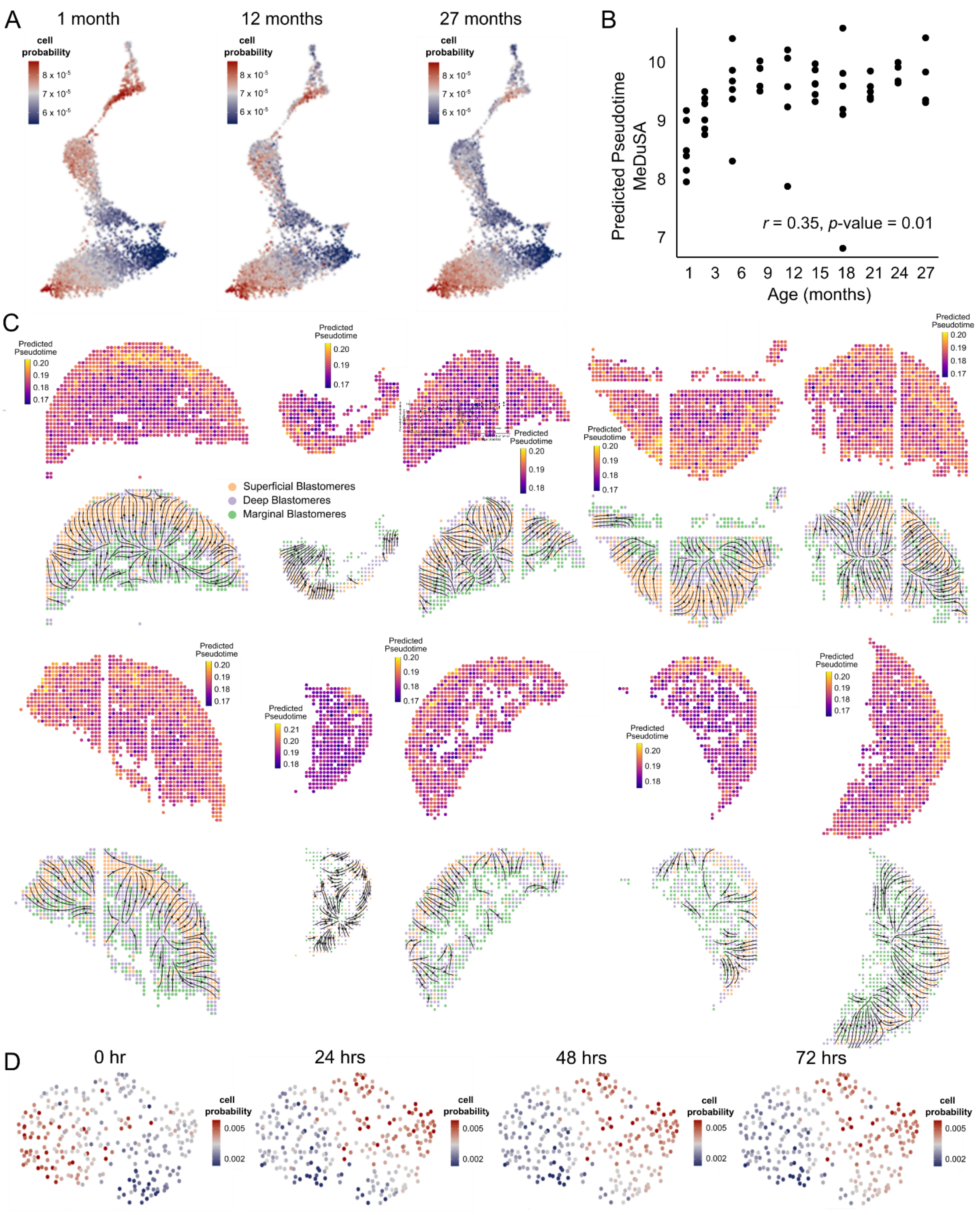


**Figure S6. Deconvolution of continuous cellular processes. (A)** Identification of age-associated changes in B-cell maturation using bulk bone marrow tissues. Single-cell abundances inferred by ConDecon for three bone marrow samples from 1-, 12-, and 27-months old mice profiled with bulk RNA-seq by the Tabula Muris Consortium[4]. **(B)** Average pseudotime inferred by MeDuSA for the B cells in each bulk sample as a function of the mice age, for bone marrow samples of 53 mice profiled with bulk RNA-seq. Pearson's correlation coefficient $r$ = 0.35, $p$-value = 0.01. **(C)** Deconvolution of spatial transcriptomic data of zebrafish embryos. Spatial tissue sections of 10 3.3 hpf zebrafish embryos profiled with Stereo-seq[5]. Each section is labeled by the average pseudotime estimated with ConDecon for the cells in each pixel (top) and the corresponding spatial cell differentiation trajectories (bottom). **(D)** UMAP representation of the single-cell ATAC-seq data of a patient-derived melanoma cell line (MM087) profiled 0, 24, 48, and 72 hours after knocking out SOX10[6]. The representation is colored by the single-cell abundances estimated with ConDecon for 4 samples from a different melanoma cell line (MM057) profiled with bulk ATAC-seq 0, 24, 48, and 72 hours after knocking out SOX10.

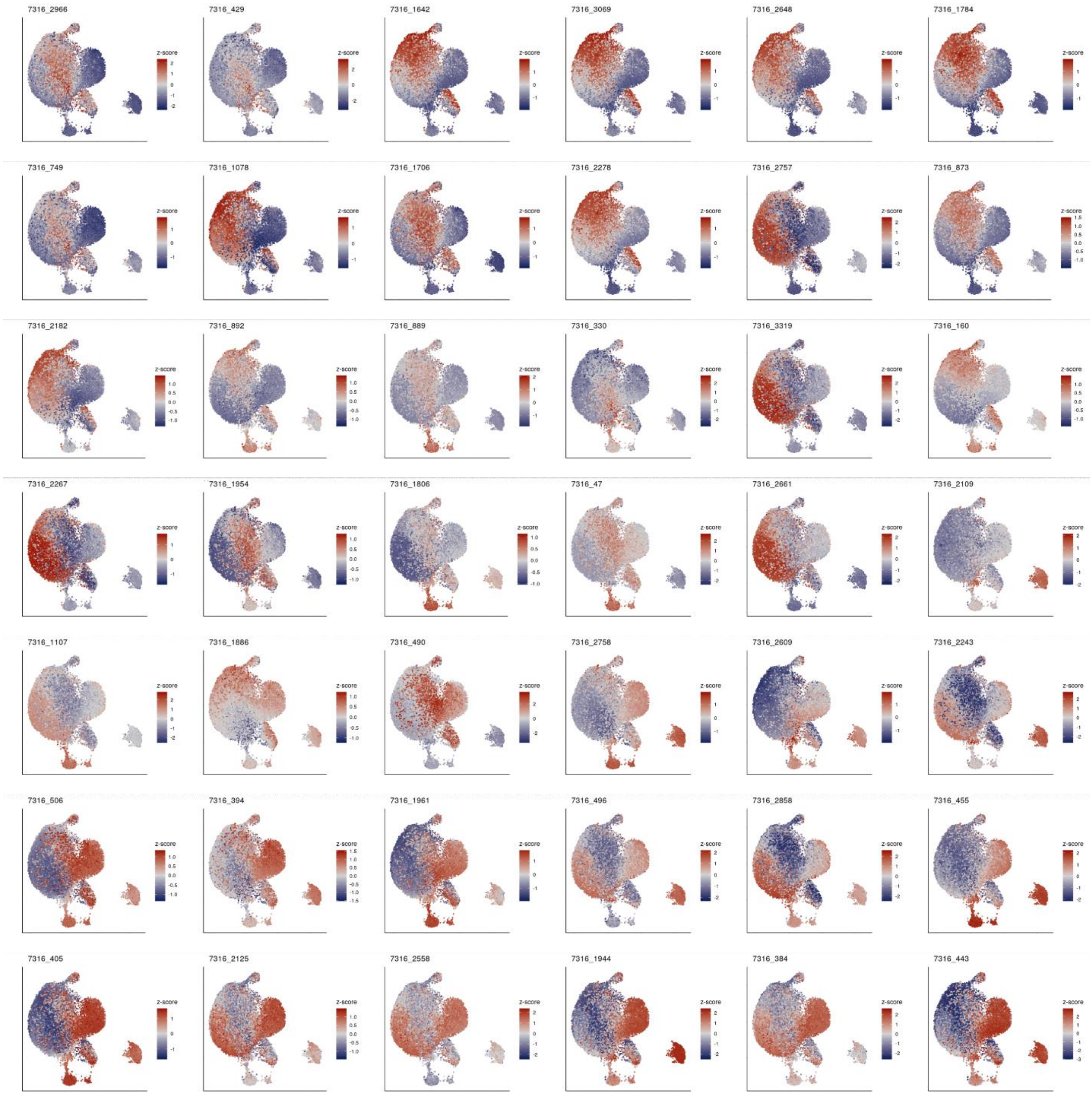


**Figure S7. Deconvolution of bulk RNA-seq data of pediatric ependymoma.** Single-cell abundance estimates computed with ConDecon for 42 pediatric ependymal tumors in the posterior fossa profiled with bulk RNA-seq. The UMAP representation of 25,349 cells from 9 posterior fossa ependymal tumors profiled with single-nucleus RNA-seq in Aubin *et al.*[7] is colored by the inferred single-cell abundances. Tumors are arranged from left to right and top to bottom according to their inferred state in the neuroepithelial-to-mesenchymal transition.

### References


1. Oetjen, K.A., Lindblad, K.E., Goswami, M., Gui, G., Dagur, P.K., Lai, C., Dillon, L.W., McCoy, J.P., and Hourigan, C.S. (2018). Human bone marrow assessment by single-cell RNA sequencing, mass cytometry, and flow cytometry. JCI Insight *3*. 10.1172/jci.insight.124928.
2. Hippen, A.A., Omran, D.K., Weber, L.M., Jung, E., Drapkin, R., Doherty, J.A., Hicks, S.C., and Greene, C.S. (2023). Performance of computational algorithms to deconvolve heterogeneous bulk ovarian tumor tissue depends on experimental factors. Genome biology *24*, 239. 10.1186/s13059-023-03077-7.
3. Newman, A.M., Steen, C.B., Liu, C.L., Gentles, A.J., Chaudhuri, A.A., Scherer, F., Khodadoust, M.S., Esfahani, M.S., Luca, B.A., Steiner, D., et al. (2019). Determining cell type abundance and expression from bulk tissues with digital cytometry. Nat Biotechnol *37*, 773-782. 10.1038/s41587-019-0114-2.
4. Tabula Muris Consortium. (2020). A single-cell transcriptomic atlas characterizes ageing tissues in the mouse. Nature *583*, 590-595. 10.1038/s41586-020-2496-1.
5. Liu, C., Li, R., Li, Y., Lin, X., Zhao, K., Liu, Q., Wang, S., Yang, X., Shi, X., Ma, Y., et al. (2022). Spatiotemporal mapping of gene expression landscapes and developmental trajectories during zebrafish embryogenesis. Dev Cell *57*, 1284-1298 e1285. 10.1016/j.devcel.2022.04.009.
6. Bravo Gonzalez-Blas, C., Minnoye, L., Papasokrati, D., Aibar, S., Hulselmans, G., Christiaens, V., Davie, K., Wouters, J., and Aerts, S. (2019). cisTopic: cis-regulatory topic modeling on single-cell ATAC-seq data. Nat Methods *16*, 397-400. 10.1038/s41592-019-0367-1.
7. Aubin, R.G., Troisi, E.C., Montelongo, J., Alghalith, A.N., Nasrallah, M.P., Santi, M., and Camara, P.G. (2022). Pro-inflammatory cytokines mediate the epithelial-to-mesenchymal-like transition of pediatric posterior fossa ependymoma. Nature communications *13*, 3936. 10.1038/s41467-022-31683-9.
