## Supplementary material for "Clustering-independent estimation of cell abundances in bulk tissues using single-cell RNA-seq data": Key Resources

### KEY RESOURCES TABLE

| REAGENT or RESOURCE | SOURCE | IDENTIFIER |
| --- | --- | --- |
| Antibodies | | |
| Anti-Human Carbonic Anhydrase IX | Novus | Cat# NB100-417SS; RRID:AB_788423 |
| Anti-Human Osteoactivin/GPNMB | R&D Systems | Cat# AF2550; RRID:AB_416615 |
| Anti-Human IBA1 | Wako | Cat# 019-19741; RRID:AB_839504 |
| Biological Samples | | |
| PFA Ependymoma FFPE tissue sections | Children's Brain Tumor Network (CBTN) | 7316-509 |
| PFA Ependymoma FFPE tissue sections | Children's Brain Tumor Network (CBTN) | 7316-490 |
| Deposited Data | | |
| Single-cell RNA-seq data of human pancreatic islands | Enge et al., 2017 | GEO: GSE81547 |
| Single-cell RNA-seq data of human pancreatic islands | Baron et al., 2016 | GEO: GSE84133 |
| Single-cell and bulk RNA-seq data of high-grade serous ovarian cancer patients | Hippen et al., 2023 | GEO: GSE217517 |
| Single-cell RNA-seq data of PBMC | 10x Genomics | <https://support.10xgenomics.com/single-cell-gene-expression/datasets/2.1.0/pbmc8k> |
| Single-cell and bulk RNA-seq and FACS data of human bone marrow cells | Oetjen et al., 2018 | GEO: GSE120446 |
| Single-cell RNA-seq data of human kidney | Han et al., 2020 | <https://figshare.com/articles/HCL_DGE_Data/7235471> |
| Bulk RNA-seq and FACS data of PBMC | Newman et al., 2019 | GEO: GSE127813; <http://cibersortx.stanford.edu> |
| Single-cell RNA-seq data from Tabula Muris Senis | Tabula Muris Consortium et al., 2020 | GEO: GSE149590 |
| Bulk RNA-seq data from Tabula Muris Senis | Tabula Muris Consortium et al., 2020 | GEO: GSE132040 |
| Single-nucleus and bulk RNA-seq data of Wilms tumors | ALSF Single-Cell Pediatric Cancer Atlas | <https://scpca.alexslemonade.org/projects/SCPCP000006> |
| Single-nucleus and bulk RNA-seq data of pediatric glioma | ALSF Single-Cell Pediatric Cancer Atlas | <https://scpca.alexslemonade.org/projects/SCPCP000010> |
| Single-nucleus RNA-seq data of ependymal tumors | Aubin et al., 2022 | GEO: GSE206578 |
| Bulk RNA-seq data of ependymal tumors | Aubin et al., 2022 | <https://portal.kidsfirstdrc.org>, project PBTA-CBTN |
| Spatial transcriptomics (Visium) data of ependymal tumors | Fu et al., 2023 | GEO: GSE195661 |
| Single-cell RNA-seq and Stereo-seq data of zebrafish embryos | Liu et al., 2022 | <https://db.cngb.org/stomics/zesta/> |
| Single-nucleus and bulk ATAC-seq data of melanoma cell lines | Bravo Gonzalez-Blas et al., 2019 | GEO: GSE114557 |
| Software and Algorithms | | |
| ConDecon (v1.0.0) | This paper | https://github.com/CamaraLab/ConDecon |
| Splatter (v1.10.1) | Zappia et al., 2017 | https://github.com/Oshlack/splatter |
| scran (v1.14.6) | Lun et al., 2016 | https://github.com/MarioniLab/scran |
| Gene expression deconvolution benchmarking pipeline | Avila-Cobos et al., 2020 | https://github.com/favilaco/deconv_benchmark |
| Seurat (v3.1.5) | Satija et al., 2015 | https://github.com/satijalab/seurat/tree/v3.1.5 |
| CIBERSORTx | Newman et al., 2019 | https://cibersortx.stanford.edu/ |
| MuSiC (v0.2.0) | Wang et al., 2019 | https://github.com/xuranw/MuSiC |
| Bisque (v1.0.5) | Jew et al., 2020 | https://github.com/cozygene/bisque |
| CPM (v 0.1.5) | Frishberg et al., 2019 | https://github.com/amitfrish/scBio |
| Monocle 3 (v0.2.3) | Cao et al., 2019 | https://github.com/cole-trapnell-lab/monocle3/tree/0.2.3 |
| Scanpy (v1.9.1) | Wolf et al., 2018 | https://github.com/scverse/scanpy/tree/1.9.1 |
| Signac (v1.1.0) | Stuart et al., 2021 | https://github.com/stuart-lab/signac/tree/1.1.0 |
| MeDuSA (v1.0) | Song et al., 2023 | https://github.com/LeonSong1995/MeDuSA |
